# Gain-controlled reconfiguration of long-range coupling explains visibility-dependent spatiotemporal neural coding dynamics

**DOI:** 10.1101/2025.09.24.677967

**Authors:** Oliver James, Yee-Joon Kim

**Author notes:** Proofs and correspondence to: Yee-Joon Kim, Center for Cognition and Sociality, Institute for Basic Science, 55 Expo-ro, Yuseong-gu, Daejeon, 34126, South Korea.

## Abstract

Numerous studies demonstrate the widespread cortical engagement in conscious and unconscious processing^1–5^, yet the mechanistic principle that reorganizes spatiotemporal neural dynamics across different levels of conscious access remains unclear. We addressed this using a personalized, connectome-constrained large-scale biophysical model of whole-brain source-imaged electroencephalography (EEG) dynamics recorded from participants who reported the orientation of a backward-masked Gabor patch at four levels of visibility. The key control parameter in the model is the global input gain (γ) to physiologically-grounded cell populations, defined operationally such that higher γ denotes weaker effective global input, which reconfigures long-range couplings and thus modulates spatiotemporal neural dynamics. Fitted γ maps showed higher gain in posterior regions and lower gain in frontal regions on visible trials, whereas the pattern was opposite on invisible trials – i.e., posterior regions received weaker and frontal regions stronger input under visibility, with the opposite under invisibility. This inversion of γ map was observed in pyramidal, VIP, and PV populations, while supplementary pyramidal, SST, and PV time constant were invariant. The visibility-dependent reversal in gain topography mirrored the observed spatiotemporal neural coding dynamics – under visibility, early occipital-to-frontal dynamic coding switched to stable coding, whereas in invisible condition, frontal-dominant stable coding emerged early with attenuated dynamic coding. Simulations generated from the fitted parameters reproduced the observed spatiotemporal neural coding dynamics, supporting a gain-control axis – with pyramidal input efficacy as the principal readout and coordinated VIP/PV modulation – that selects between locally integrating and globally broadcasting regimes as visibility changes. Our results provide a bottom-up mechanistic account of differential perceptual processing depending on the degree of the stimulus awareness, offering insights into potentially resolving ongoing debates about theories of consciousness.

## Main

Conscious access, the process by which subjective experiences become available for explicit report, is a fundamental aspect of human perception^6,7^. Recent neuroimaging works near perceptual threshold have revealed that conscious and unconscious processing engage widely distributed cortical – and, at times, subcortical – networks^1–5^. One EEG study applying the temporal generalization techniques to EEG activity patterns has demonstrated that an early time-changing activity pattern (dynamic neural code) in posterior area shifts to a later sustained activity pattern (stable neural code) during conscious access, while the stable neural code emerges early in frontal area under low subjective visibility^5^. Another EEG study has shown that a stimulus close to the threshold yields a mixture of high and low conscious access trials, producing a burst in the trial-to-trial variability of conscious access revealing bifurcation dynamics around 250 – 300 ms that predict conscious access even without report^1^. Yet, despite this wealth of findings, the conclusions usually remain phenomenological – often phrased as consistency with a favored theory of consciousness – without identifying a precise mechanistic principle that selects among observed spatiotemporal neural dynamics at the boundary between aware and unaware perception.

Here, to gain mechanistic insights into these visibility-dependent spatiotemporal neural dynamics, we therefore developed a personalized, connectome-constrained biophysical whole-brain model that places a physiologically-grounded canonical mesoscopic microcircuit inside in each of 68 cortical regions defined by a standard atlas connected according to each participant’s anatomical connectivity (MRI/DTI) (Fig.1a. See Methods for details). This approach is inspired by, but not identical to, prior large-scale dynamical models that couple single-population excitatory and inhibitory units by the human and non-human primate connectome^8–10^. In contrast to those minimal node models, we instantiate a five-population canonical circuit at each region – pyramidal (PY), parvalbumin-expressing (PV), vasoactive-intestinal-peptide (VIP), somatostatin (SST), and a supplementary pyramidal pool (sPY) – because these populations and their qualitative roles are well established in human and animal work^11,12^ (Fig. 1b). Using a state-space framework, we fit the model to the source-imaged EEG signals, estimating key global input gain parameters (γ) that modulate long-range interactions between regions (Fig. 1c). This approach links the mesoscopic circuit properties to the observed macroscopic EEG brain dynamics, providing a multi-scale perspective on conscious access. Long-range coupling is personalized by each participant’s structural connectome derived from MRI/DTI, and a compact set of effective input gains is estimated from the data for each visibility condition.

**Fig. 1.**
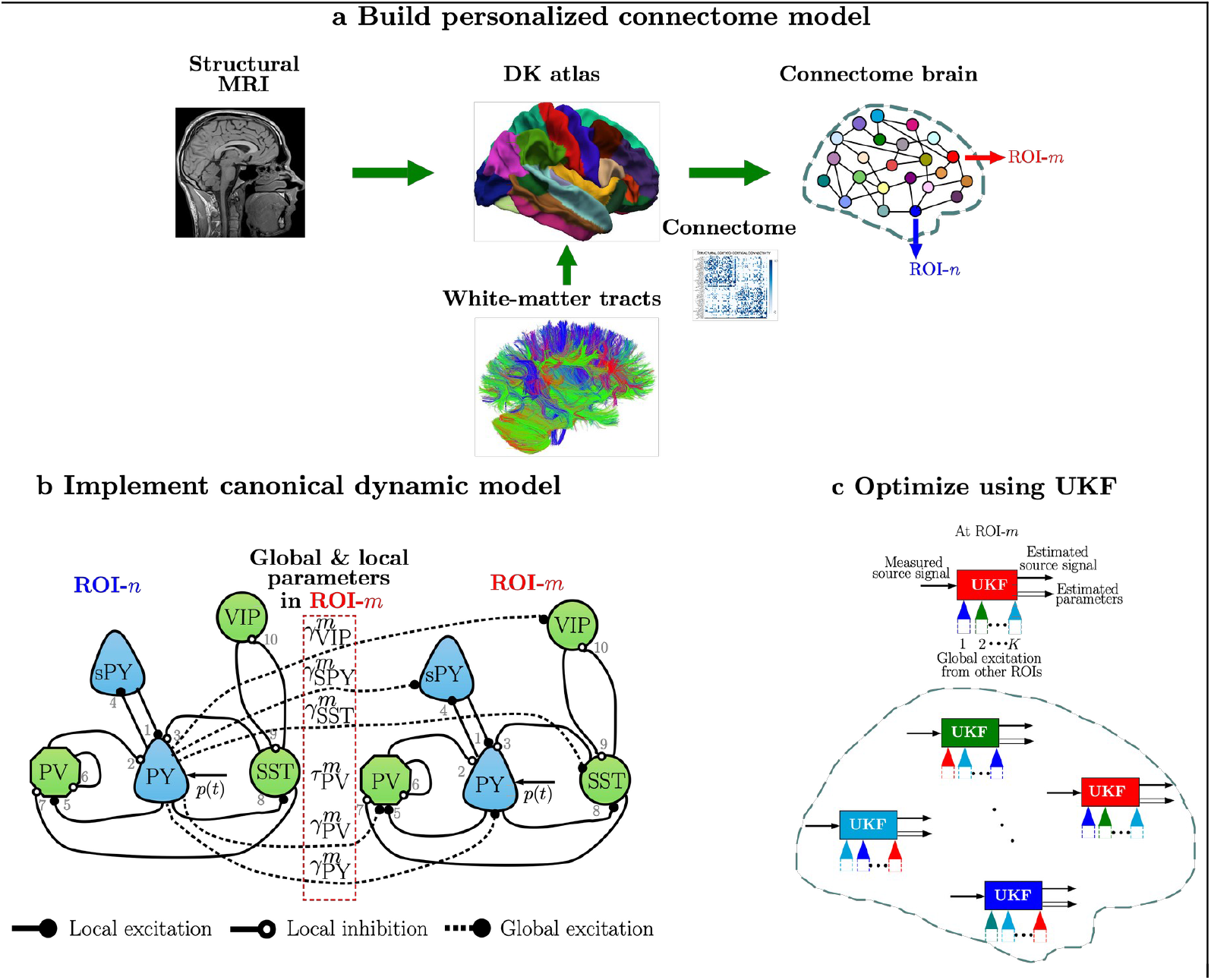
Large-scale dynamic modeling of personalized connectome and parameter estimation. **a**. Workflow for constructing a personalized connectome model using structural MRI and diffusion tensor imaging (DTI) data. This approach integrates individual anatomical information to create a participant-specific brain network model. **b**. Integration of a canonical dynamic circuit model within the nodes of the personalized connectome. The local and global parameters of the canonical model to be estimated from source data are highlighted in the red dashed box. **c**. Unscented Kalman filter (UKF) implementation in each node to estimate regional local and global parameters. The UKF, a nonlinear state estimation technique, allows for simultaneous tracking of neural states and parameter values based on observed brain activity.

This approach allows us to investigate whether the visibility-dependent differential spatiotemporal neural coding dynamics arise from either the properties of local population activity within each brain region or the mutual influences that these populations exert on each other, as mediated by their interconnecting anatomical connections. We hypothesized that near-threshold perception would elicit distinct changes in both behavior and brain activity despite minimal stimulus difference as shown in previous studies^1,5^ and that our model’s parameters would capture distinct network regimes corresponding to conscious vs. unconscious states. By combining source-imaged EEG with computational modeling, we aimed to reveal the spatiotemporal neural dynamics governing the tipping point of awareness and to place mechanistic constraints on current theories of consciousness.

### Behavioral indicators of conscious access levels

As our goal was to explain, in bottom-up terms, how a small set of circuit parameters can govern the pattern of spatiotemporal neural dynamics as stimulus visibility varied, we used a well-established backward-masked visual orientation task^13^ combined with source-imaged EEG to replicate previously published research findings^5^. We analyzed EEG data from 27 participants while they viewed a target grating. Using a staircase procedure, we titrated mask contrast level such that two critical conditions – subjectively visible (SV) and subjectively invisible (SI) – had nearly identical stimulus contrast^14^. Alongside these, we included clearly visible (OV, objectively visible) and completely invisible (OI, objectively invisible) conditions to span the full range of conscious access (Fig. 2a). This taxonomy allowed us to compare neural responses when physical stimulation was closely matched yet subjective access differed, and conversely, when access was similar despite differences in objective discriminability.

**Fig. 2.**
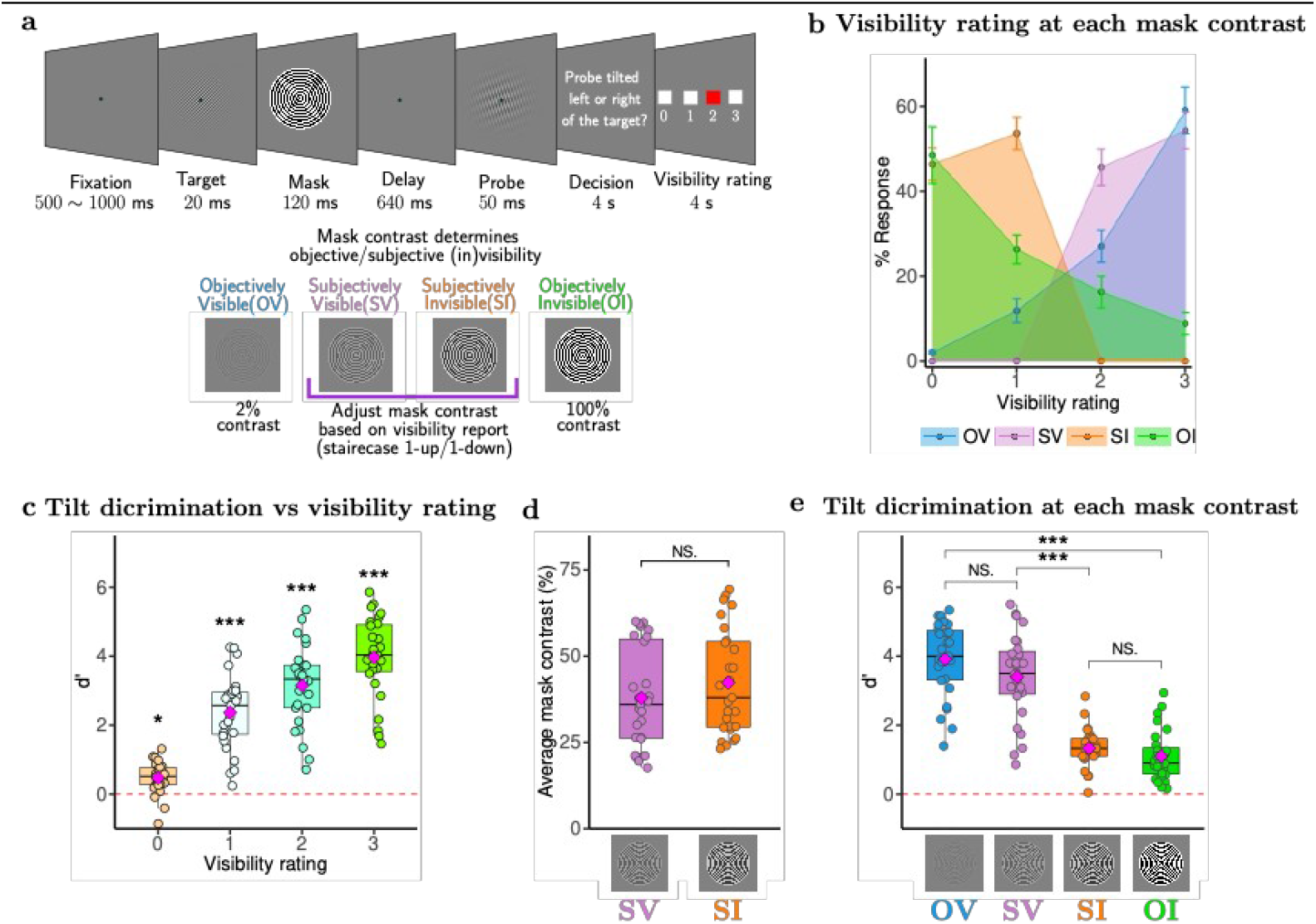
EEG experimental paradigm and behavior performances (N = 27). **a**. Participants were instructed to maintain the orientation of a masked Gabor patch to compared it to a subsequent probe. Participants reported whether the probe orientation was rotated clockwise or counterclockwise relative to the target orientation. At each trial, participants reported the target visibility with a four-point scale. In the OV and OI condition, the mask contrasts were fixed to 2% and 100%, respectively. The mask contrasts for subjective conditions were determined based on the visibility rating using 1up-1down staircase procedure. **b**. The proportion of target visibility reports for each level of mask contrast confirms that participants adequately used the visibility ratings. Error bars indicate the SEM across participants. **c**. Tilt discrimination performance *d*’ as a function of visibility rating. **d**. Mask contrast in staircase trials. The mean (diamond shape) mask contrast between subjective conditions is insignificant, indicating physically similar contrast levels. **e**. Replicating Bae *et al*. (2025), forced-choice tilt discrimination index (*d*’) for the four conditions reveals significant differences between visibility (OV/SV) and invisibility (SI/OI) conditions.

An experimental trial starts with a central fixation. After 500 ∼ 1000 ms, a target Gabor patch appeared for 20 ms followed by a backward mask with a variable contrast (Fig. 2a). On every trial, participants report whether the probe orientation was tilted left or right relative to the target orientation. We also asked participants to indicate their visibility rating from in a four-point visibility scale (0, completely unseen; 3, clearly seen). The visibility report percentages (Fig. 2b) indicate that participants effectively utilized the subjective visibility ratings, demonstrating their understanding of the rating scale. We next assessed participants’ performance using the discriminability index (*d’*) that determined the ability of the participants to correctly report the orientation of the masked Gabor patch (the target) and compared it to the subsequent probe. Across all trials, we observed that discrimination performance improved monotonically with reported visibility: d′ increased steeply from 0 (fully unseen) to 3 (fully seen) on the subjective scale (Fig. 2c). This pattern also reveals that as participants reported higher levels of visibility, their ability to discriminate the target stimulus improved markedly. Remarkably, even when participants reported that the target was invisible (scale 0), they still demonstrated a weak but significant behavioral performance (*t(26) = 5*.*29, p < 0*.*001*). To probe the near-threshold regime of conscious access where subjective experience teeters between visible and invisible despite minimal differences in stimulus input, we classified trials by manipulating mask contrast. Objectively visible (OV) and invisible (OI) conditions used fixed contrasts of 2% and 100% respectively. Unlike OV and OI conditions with the fixed mask contrast levels, the 1up-1down staircase procedure ensured that the mask contrasts in SV and SI conditions were virtually identical (Fig. 2d; SV: 38 ± 2.8 %, SI: 42 ± 12.9 %. *t(26) = 1*.*09, p = 0*.*282*). Despite this, participants’ ability to discriminate the target’s orientation differed dramatically between SV and SI. The behavioral accuracy in subjectively visible trials was significantly higher than in subjectively invisible trials (Fig. 2e; *t(26) = 7*.*88, p*<0.001), even though physical stimuli were nearly identical. This striking divergence suggests that internal network state, rather than stimulus strength, determined the behavioral outcome, implicating that variations in stimulus awareness are likely driven by top-down modulatory processes when bottom-up sensory input remains constant^5,15–17^. Indeed, theories emphasizing access-stage of consciousness predict such a scenario, where constant sensory input yields variable awareness due to fluctuating brain states^18^. The *d’* (Fig. 2e) was high in both visible conditions: OV (3.91 ± 0.21) and SV (3.40 ± 0.24) with their mean difference remain insignificant (*t(26) = 1*.*61, p = 0*.*114*). Also, participants performed above chance (though with low d′) even in “invisible” conditions (SI: 1.33 ± 0.11 and OI: 1.01 ± 0.14), indicating residual visual processing of the masked stimuli despite the absence of their visibility. We noted that *d’* was significantly different across the conscious access levels (Linear mixed model, *F(3,78) = 120*.*6, p < 0*.*001, Cohen’s f*^*2*^*=1*.*73*).

### Visibility-dependent spatiotemporal neural coding dynamics

Next, we performed a temporal generalization (TG) analysis on these four datasets in order to investigate the spatiotemporal neural coding profiles of target orientation as a function of conscious access at the scalp and in source-localized regions of interest (ROIs) that tile occipital, parietal, temporal, and frontal cortices (See Methods for details). Briefly, to discriminate all possible pairs of target orientations, we trained linear discriminant analysis (LDA) classifier on OV stimulus-evoked EEG activity patterns for a pair of target orientations at a particular time point and the generalized it to the EEG signals over the entire period. We iterated this procedure until the classifier trained at every time point was generalized to the EEG signals obtained at every time point, to make a two-dimensional TG matrix of decoding accuracies (5-fold cross-validation). A comparable mean trial count in each condition is ensured to reduce the potential classifier biases (Extended Data Fig. 1). As the stimulus was most visible in the OV condition, we applied this decoder trained on the OV condition to other conditions. We used the AUC as a measure of the accuracy of decoding the backward-masked orientation.

#### Sensor-level temporal generalization

We performed a TG analysis separately for the three electrode clusters (37 posterior, 37 central, and 36 anterior electrodes) to examine the role of each region in stimulus information processing (Three rows of two-dimensional TG maps in Fig. 3). As TG methods reveal whether the neural code that supports above-chance decoding of orientation percepts is stable or dynamically evolving over time, we also summarized these patterns of two-dimensional modulations of decoding accuracies in the TG map using a time-resolved dynamic and stable index ^5^ (Line plots below each TG map in Fig. 3. See Methods for the details). Stable neural code refers to a situation where the decoder trained at a particular time point can reflect the presence of orientation information over a different period, resulting in the rectangle-shaped decoding accuracy modulations in the TG map. Dynamic neural code refers to a situation where the decoder trained at a particular time point can reflect the presence of orientation information only at the matched time point, resulting in the significant decoding accuracy modulations along the diagonal axis in the TG map. Briefly, the dynamic index is the proportion of the off-diagonal elements that were significantly smaller than the corresponding on-diagonal elements. The stable index is the proportion of the off-diagonal elements that were significantly greater than 0.5 and not significantly smaller than the corresponding on-diagonal elements.

**Fig. 3.**
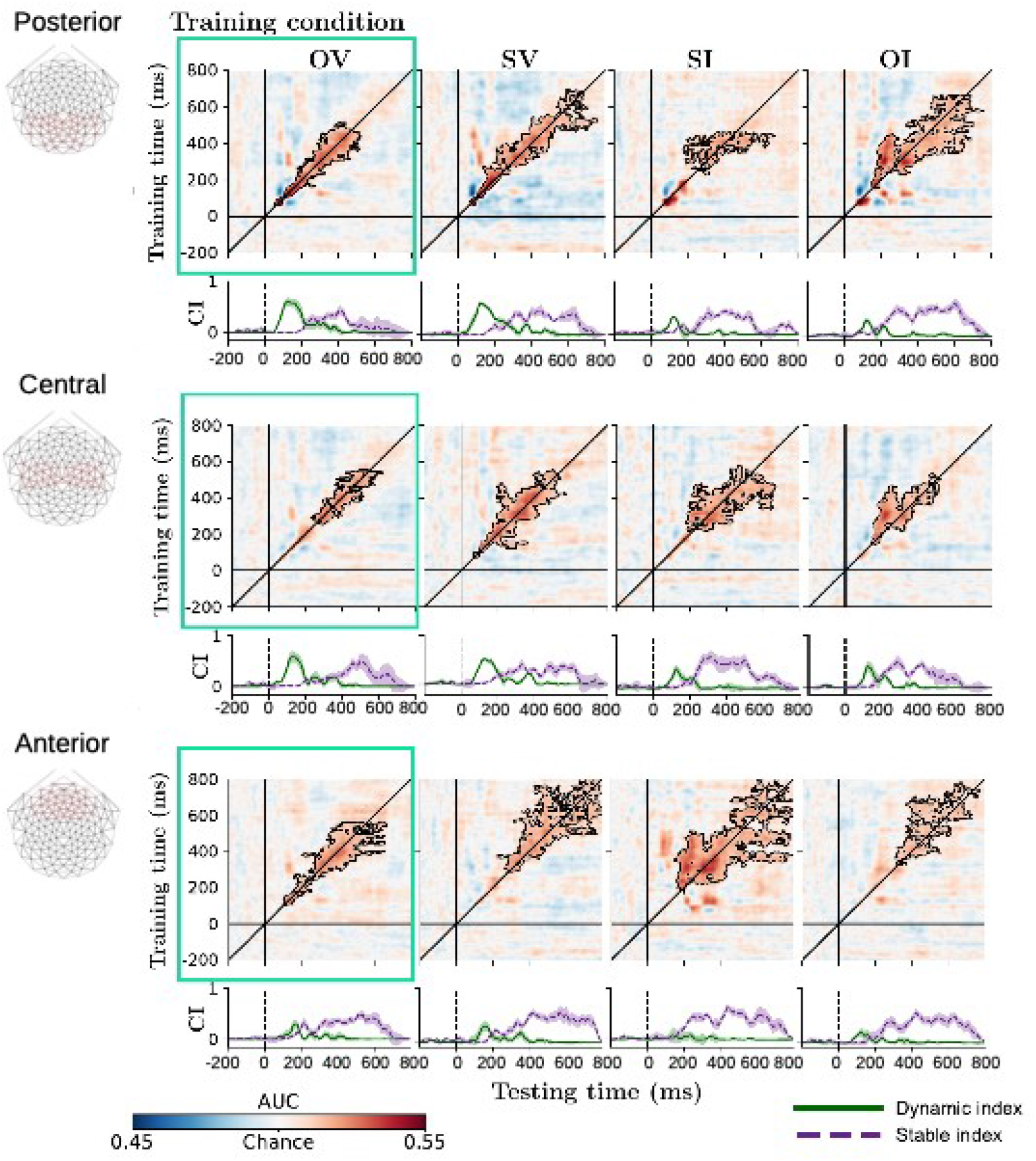
Sensor-level spatiotemporal neural dynamics as a function of visibility (N = 27). TG maps and dynamic/stable index of each dataset for the posterior, central, and anterior region (see electrode maps on the left). Colors represent the decoding accuracy (AUC). Black contours indicate clusters of significant decoding accuracies above chance (*p < 0*.*05*, based on cluster extent). The x- and y-axis indicate the time of testing and training sets after the stimulus onset, respectively. Green and purple line plots under each TG map show the dynamic and stable index summarizing the temporal coding of TG, respectively. The x-axis shows the time after stimulus onset. The y-axis shows the magnitude of stable and dynamic index. Shaded areas represent ±1 bootstrapped standard error. CI: Neural code index.

In Figure 3, the four columns, from left to right, represent OV, SV, SI, and OI and the three rows from top to bottom represent the TG matrices and their time-resolved dynamic and stable index for the posterior, central and anterior electrodes, respectively. This table summarizes how the presence of orientation information is reflected in different brain regions over time as conscious access to the identical stimulus varied. The TG maps of all three brain regions showed significant time clusters where the decoding accuracy was significantly higher than chance, as indicated by the black contours. As these results replicated previous perceptual integration study that has demonstrated the visibility-dependent spatiotemporal neural coding dynamics^5^, we will summarize them in terms of distinct neural coding patterns rather than detailing the coding dynamics in each panel of the table of TG matrices. In OV and SV, posterior electrodes exhibited a pronounced early dynamic neural code that later transitioned to stable neural code (The 1^st^ and 2^nd^ columns of Fig. 3). In SI and OI, early dynamic neural codes were attenuated in all three brain areas, whereas stable neural codes emerged earlier and more broadly in anterior regions (The 3^rd^ and 4^th^ columns of Fig. 3). The distinct neural coding of posterior and anterior electrodes according to the level of conscious access and the dominant involvement of anterior regions in stimulus processing under low visibility suggest the need for a mechanism that selectively re-allocates long-range influence across brain areas as visibility changes.

#### Source-level temporal generalization

Source space data were obtained by projecting the sensor data on to the inverse matrix (15,000 cortical surface vertices x 128 electrodes), calculated using dynamical statistical parametric mapping (Methods). The cortical source vertices were organized based on the DK atlas that assigns vertices of 68 regions of interest ROI (34 per hemisphere). The TG analyses on 16 separate bilateral ROIs (combining the responses in the right and left hemispheres. See Methods) revealed the same qualitative organization of spatiotemporal neural codes as observed at the sensor-level. Figure 4 shows TG maps and their corresponding coding indices in four representative ROIs^13^ (Lingual gyrus L, Superior Parietal cortex SP, Inferior Temporal cortex IT, and Rostral Middle Frontal cortex RMF. See Fig. 6a for the remaining 12 ROIs). TG maps for all four ROIs showed significant time clusters as indicated by black contours lines. When target orientation was visible in OV and SV condition, these ROIs almost concurrently showed the significant decoding accuracy modulations in the TG maps, which were effectively captured by dynamic neural codes during a period from ∼75 ms to 350 ms after the stimulus onset in L, SP and IT regions while the RMF displayed brief dynamic neural codes over the post-stimulus period from ∼80 ms to 200 ms (Green line plots in the 1^st^ and 2^nd^ columns of Fig. 4). Around 200 ms after stimulus onset, we observed stable neural codes in both OV and SV conditions (Purple line plots in the 1^st^ and 2^nd^ columns of Fig. 4).

**Fig. 4.**
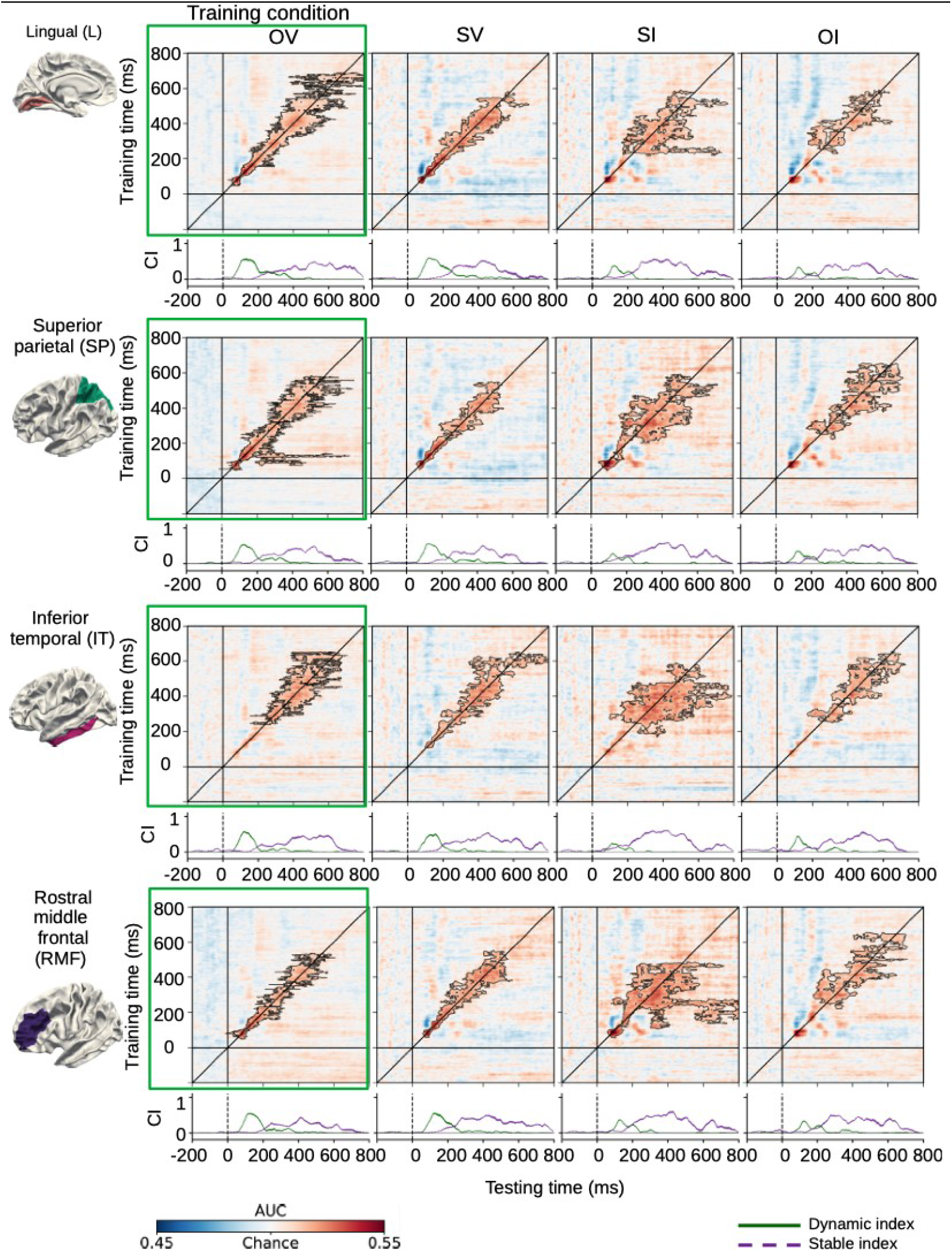
Source-level spatiotemporal neural dynamics as a function of visibility (N = 21). TG maps and dynamic/stable index of each dataset for four representative source regions: Lingual gyrus L, Superior Parietal cortex SP, Inferior Temporal cortex IT, and Rostral Middle Frontal cortex RMF (see ROIs on the left). Colors represent the decoding accuracy (AUC). Black contours indicate clusters of significant decoding accuracies above chance (*p < 0*.*05*, based on cluster extent). The x- and y-axis indicate the time of testing and training sets after the stimulus onset, respectively. Green and purple line plots under each TG map show the dynamic and stable index summarizing the temporal coding of TG, respectively. The x-axis shows the time after stimulus onset. The y-axis shows the magnitude of stable and dynamic index. Shaded areas represent ±1 bootstrapped standard error. CI: Neural code index.

When Gabor patch was not detected with clear sense of its orientation, dynamic neural code index was much reduced compared with when the target orientation was visible (Green line plots in the 3^rd^ and 4^th^ columns of Fig. 4). However, despite the fact that the stimuli were nearly identical in the SO and SI conditions (Fig. 1d), the TG maps were dramatically different between the two conditions – TG maps in the SO condition showed that dynamically evolving activity patterns dominated, producing narrow diagonal-shaped significant time clusters (The 2^nd^ column of Fig. 4; OV/SV conditions in Extended Data Fig. 2), while TG maps in the SI condition showed that sustained activity patterns dominated, producing wide rectangle-shaped off-diagonal significant time clusters (The 3^rd^ column of Fig. 4; SI/OI conditions in Extended Data Fig. 2). Importantly, in the SI condition, stable neural representation of the target orientation emerged earlier in frontal cortex than in visual cortex (Purple line plots in the 3^rd^ column of Fig. 4). This suggests that when the stimulus failed to trigger a strong bottom-up sweep, the frontal regions were nevertheless activated (perhaps by prior expectation) and sent top-down signals to sensory regions in an attempt to refine or rescue the weak stimulus representation^19^. Indeed, the dynamic coding in SI trials was relatively weak and short-lived, especially in frontal cortex, implying a failure to ignite a prolonged global broadcast. Taken together, these results point to two distinct regimes of cortical processing near the conscious threshold: one dominated by bottom-up activation spreading from sensory to frontal areas when the stimulus is successfully perceived., and one dominated by early top-down feedback from frontal areas to sensory cortex when the stimulus remains unconscious.

### Global input gain parameter reconfiguration underlying neural coding dynamics

TG maps showed the visibility-dependent spatiotemporal dynamics in both sensor and source space – posterior-led early dynamic code followed by later stable code under visibility, versus earlier frontal stable code under invisibility. This double dissociation in the organization of the code geometry led us to ask whether a small set of global input gains in our biophysical model can account for this redistribution of long-range influence (Fig. 1) and thereby explain the visibility-dependent TG patterns mechanistically.

We estimated latent states and circuit parameters concurrently in all ROIs by using a state-space approach that respects signal delays introduced by long-range connections. This concurrent optimization procedure mimics the parallel stimulus processing of interconnected brain regions, allowing model parameters to be estimated simultaneously across all ROIs. As algorithmic details are provided in Methods, here we emphasize the modeling logic. First, we fit our large-scale cortical connectome model to source-imaged EEG data separately for each visibility condition by running all unscented Kalman filters^20–22^ (UKFs) concurrently in 68 ROIs (Fig. 1c). This allowed γ parameters to differ by visibility condition while keeping the local circuit connectivity strengths (Black solid line in Fig. 1b) in each ROI fixed to biophysically plausible values based on previous studies^12,23^ (Extended data table 1). For each condition, one UKF per ROI yielded converged values and convergence times for six model parameters such as one local PV time constant (*τ* _*PV*_) that is believed to stabilize sensory representation^24,25^ and five global input gain parameters (*γ* _*PY*_, *γ* _*PV*_, *γ*_*VIP*_, *γ* _*SST*_ and *γ* _*sPY*_) that are inversely correlated with the strengths of the incoming neural signals from all other ROIs: high and low global parameter values indicate weak and strong input signals that each ROI receives from other ROIs, respectively (Black dotted lines connecting the two ROIs in Fig. 1b). Second, we used the fitted parameters to run forward simulations from the empirical inputs, generating synthetic time series that we analyzed with the same TG pipeline used for the data. This two-step loop – fit then simulate – tests whether a small set of gain parameters can reproduce the observed code geometry and whether condition differences appear as a reconfiguration of global input gain.

Our vertex-wise fitting procedure allowed us to visualize the complete spatial distribution of FDR-corrected average global input gain parameter across the entire cortical surface as a function of visibility (See Methods for details). The spatial topology of *γ* _*PY*_ inverted between visible (OV/SV) and invisible (SI/OI) conditions (Fig. 5a). Under visibility, occipital, parietal, and temporal cortices exhibited higher *γ* _*PY*_ (weaker effective global input) and frontal cortices lower *γ* _*PY*_ (stronger input). Under invisibility the pattern reversed: frontal *γ* _*PY*_ increased and occipital, parietal, and temporal *γ* _*PY*_ decreased. We also found that *γ*_*VIP*_ and *γ* _*PV*_ in the model (affecting inhibitory cell types) displayed similar spatial trends across conditions (Extended Data Fig. 3 & 4), suggesting a broad reconfiguration of cortical circuitry between conscious and unconscious states. These gain topographies provide an integrated view of how the system transitions between locally integrating and globally broadcasting regimes as conscious access changes. By contrast, *γ* _*SST*_ (Extended Data Fig. 5), *γ* _*sPY*_ (Extended Data Fig. 6), and *τ* _*PV*_ showed no reliable condition-dependent differences.

**Fig. 5.**
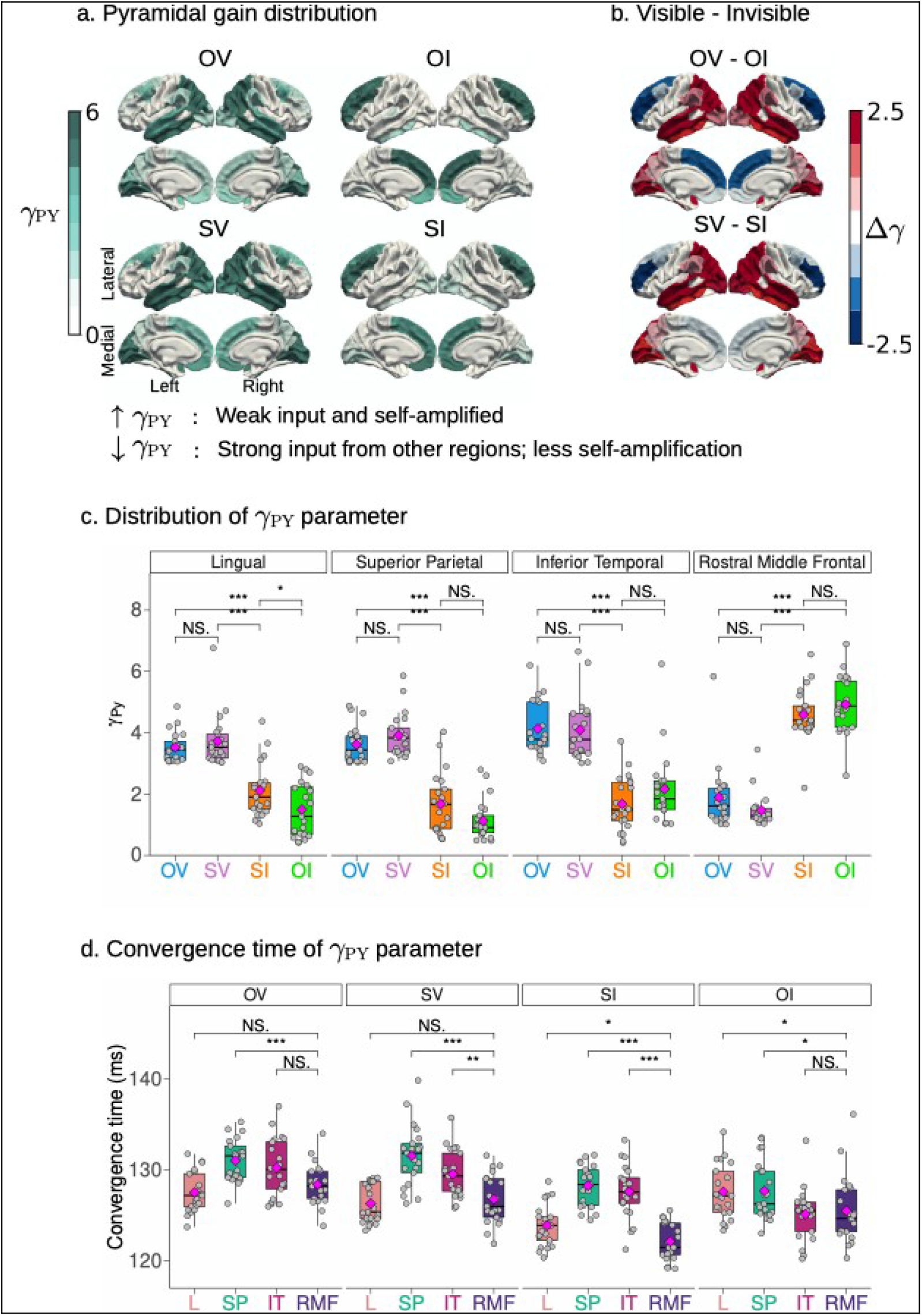
Comprehensive surface maps of pyramidal cell global input gain (*γ* _*PY*_) distribution reveals differential engagement during varying levels of conscious access. **a**. Vertex-level surface plots of *γ* _*PY*_ for OV, SV, SI, and OI conditions. Higher *γ* _*PY*_ values (dark green) indicate weaker global input from other cortical areas. OV/SV conditions show higher *γ* _*PY*_ in visual, parietal, and temporal cortices, with lower values in prefrontal cortex. SI/OI conditions indicate the opposite pattern, with lower *γ* _*PY*_ in sensory areas and higher values in frontal regions. **b**. Vertex-wise difference in the *γ* _*PY*_ values between objective (OV/OI) and subjective (SV/SI) conditions. Red colors indicate higher *γ* _*PY*_ values in visible condition (SV > SI, OV > OI), while blue areas show higher *γ* _*PY*_ in invisible conditions (SI > SV, OI > OV). Despite similar stimulus strengths, SV/SI comparisons reveal striking spatial differences. SV condition shows increased *γ* _*PY*_ in visual, parietal, and temporal cortices. SI condition exhibits elevated *γ* _*PY*_ in frontal cortex. **c**. The averaged *γ* _*PY*_ (mean of left and right hemisphere values) in four representative ROIs. The averaged *γ* _*PY*_ differences (OV – OI and SV – SI) remained statistically significant in all four ROIs (L: OV: 3.52 ± 0.49, OI: 1.49 ± 0.88; *p < 0*.*001*; SV 3.73 ± 0.84; SI: 2.11 ± 0.87, *p < 0*.*001*; SP: OV: 3.61 ± 0.59; OI: 1.12 ± 0.66; *p<0*.*001*; SV: 3.92 ± 0.71; SI: 1.68 ± 0.98; *p < 0*.*001*; IT: OV: 4.13 ± 0.84; OI: 2.16 ± 1.18; *p < 0*.*001*; SV: 4.09 ± 0.97; SI: 1.59 ± 0.88; *p < 0*.*001*; RMF: OV: 1.89 ± 1.03; OI: 4.92 ± 0.95; *p < 0*.*001*; SV: 1.47 ± 0.54; SI: 4.52 ± 0.83; *p < 0*.*001*), with a weak but significant mean difference in L between SI/OI (SI: 2.11 ± 0.87; OI: 1.49 ± 0.88; *p < 0*.*05*). **d**. Convergence time for *γ* _*PY*_ at various cortical locations weakly co-vary with *γ* _*PY*_ (Extended Data Fig. 9).

To clarify the condition-dependent spatial gradients of *γ* _*PY*_, we summarized the visible – invisible difference in γ maps (Δγ) (Fig. 5b) and the objective – subjective differences in γ maps (Δγ) (Extended Data Fig. 2). As the reversal in gain topography evident in Figure 5b is reminiscent of distinct neural code geometry observed in Figure 4, we related gain parameters to TG patterns. Regions with higher *γ* _*PY*_ under invisibility – notably frontal ROIs – exhibited earlier and broader stable code (larger stable index), whereas posterior regions with lower *γ* _*PY*_ under visibility expressed stronger early dynamic code (larger dynamic index). These distinct patterns of cortical engagement under different visibility were evaluated by comparing the global input gain (averaged across all vertices) between conditions within four representative ROIs (Fig. 5c). Especially, between the two near-threshold conditions (SV vs. SI), global input gains of pyramidal neuron were significantly different in four representative ROIs despite nearly identical stimulus intensities (SV vs. SI: *p* < 0.001 for each of lingual gyrus, superior parietal, inferior temporal, and frontal ROI; Fig. 5c). This confirms that the brain operates in a different network mode depending on subjective visibility. Specifically, this dissociation suggests that subjective awareness may be associated with enhanced processing in sensory areas, whereas the absence of awareness involves greater frontal lobe activity, possibly reflecting altered information flow although we cannot exclude the possibility that some residual weak stimulus information may be rescued by frontal activation due to trial-to-trial variability of conscious access in SI condition ^1,19^. The spatial distribution *γ*_*VIP*_ and *γ* _*PV*_ show patterns similar to that of *γ* _*PY*_ but not the parameters *γ* _*SST*_ and *γ* _*sPY*_ (Extended Data Fig. 7a – 7d) in the four ROIs. The local PV time-constant parameter *τ* _*PV*_ does not show significant differences among the ROIs (Extended Data Fig. 7e) implying consistent temporal processing characteristics regardless of their location. While correlational, these associations demonstrate that the same spatial gradients that organize γ reconfiguration also organize the redistribution between dynamic and stable codes across visibility.

Concurrent UKF fits also yielded convergence time distributions for the six parameters across all ROIs (Fig. 5d and Extended Data Fig. 8). Convergence times (averaged across hemispheres) showed complementary structure: In SI and OI, high *γ* _*PY*_ in RMF converged earlier than those in L, SP, and IT, with weaker or inconsistent spatial gradients of convergence times in OV and SV (Fig. 5d). This dissociation suggests that when conscious access fails, frontal circuits rapidly settle into a top-down stable regime, while sensory regions converge more slowly under weak or degraded feedforward drive. Conversely, the lack of convergence time differences in visible conditions may emphasizes successful integration within posterior cortices. Such condition-dependent convergence dynamics align with the global neural workspace theory (GNWT) framework, in which frontal stabilization without posterior ignition leads to unconscious processing. Conversely, the lack of convergence time differences in visible conditions is consistent with integrated information theory (IIT), emphasizing successful integration within posterior cortices. Taken together, these results suggest that GNWT and IIT may capture complementary aspects of conscious access, with convergence time serving as a dynamical marker of whether perception is feedforward-driven (visible) or feedback-driven (invisible). The correlation pattern between convergence time and *γ* _*PY*_ changes gradually from posterior to frontal areas (Extended Data Fig. 9) suggests a potential link between local circuit properties and their coding dynamics.

Forward simulations using the fitted parameters reproduced the key qualitative features of the TG maps across visibility levels (Fig. 6b): a posterior early dynamic code followed by stable code under visibility, and earlier frontal stable code with attenuated dynamic code under low access. Because the synthetic data were processed identically to the empirical data, this agreement supports the interpretation that a low-dimensional reconfiguration of global input gains across PY/VIP/PV is sufficient to explain the reorganizations of the observed code geometry in a connectome-constrained, canonical mesoscopic model.

**Fig. 6.**
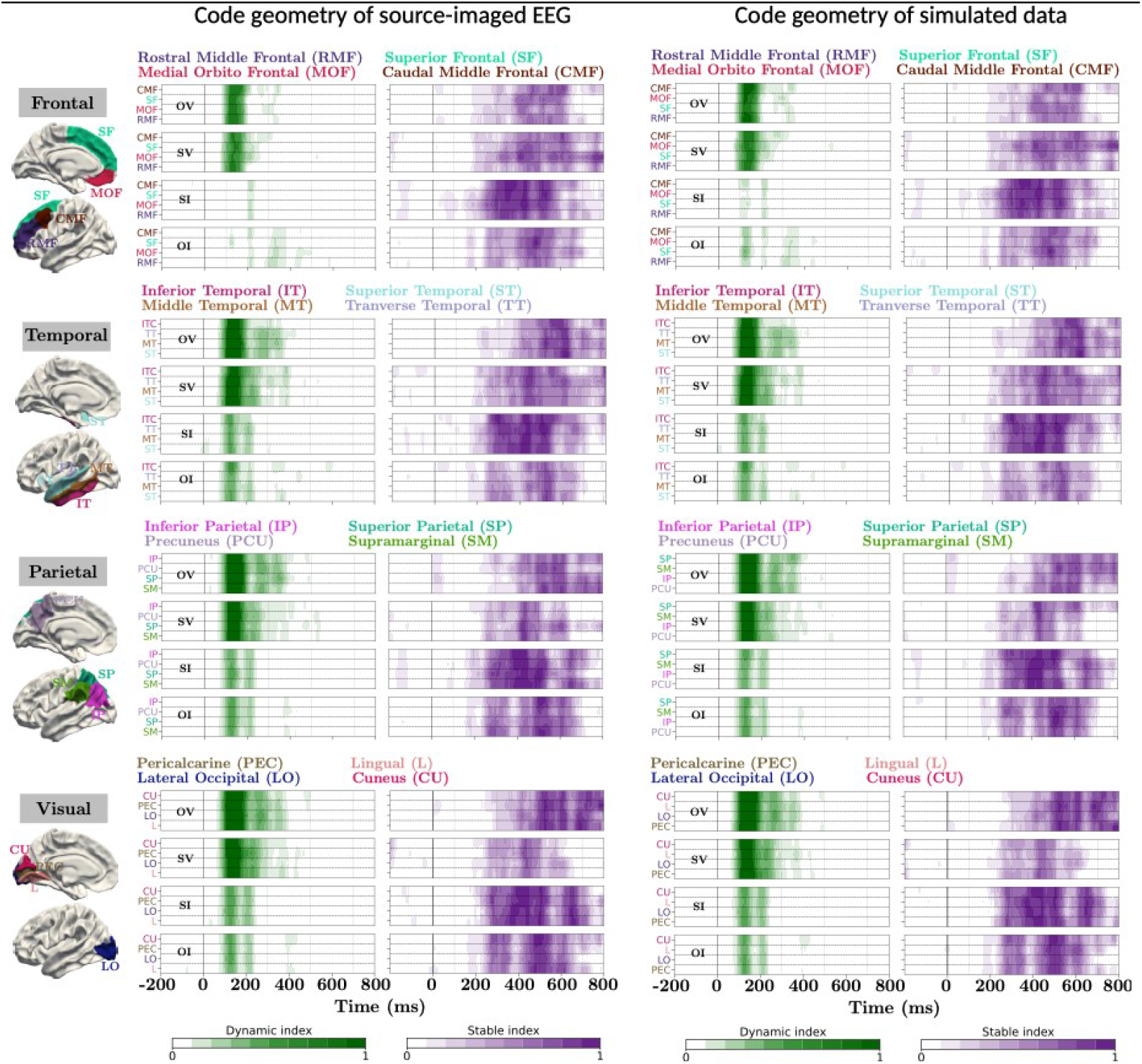
Dynamic/stable indices of source-imaged EEG data and simulated data in 16 ROIs. **a**. Dynamic/stable indices of source-imaged EEG data for each visibility condition in 16 ROIs The regions include – **Frontal:** Rostral middle frontal (RMF), Superior frontal (SF), and Caudal middle frontal (CMF), Medial orbitofrontal (MOF); **Temporal:** Inferior temporal cortex (ITC), Superior temporal (ST), Transverse temporal (TT), and Middle temporal (MT); **Parietal:** Superior parietal (SP), Inferior parietal (IP), Supramarginal (SM), and Precuneus (PCU); **Occipital:** Lingual (L), Pericalcarine (PEC), Lateral occipital (LO), and Cuneus (CU). The contrasting spatial patterns across conditions suggest a differential involvement in dynamic and stable information processing depending on anatomical location and functional specialization. **b**. Dynamic/stable indices of simulated data for each visibility condition in 16 ROIs. The close correspondence in TG maps between simulated data and actual data validates the reliability of our model’s data fitting procedure and suggests that the estimated parameters are accurate. This comparison provides strong support for the robustness of the model and validity of the observed neural coding patterns across different brain regions. The green and purple colormaps indicate dynamic and stable indices, respectively.

## Discussion

We set out to provide a mechanistic account beyond a phenomenological description of how visibility reorganizes spatiotemporal neural coding dynamics. By embedding a canonical five-population circuit inside a participant-specific connectome and fitting global input gains under each visibility condition, we showed that a low-dimensional reconfiguration of global input gains across pyramidal, VIP and PV populations explained the observed spatiotemporal coding dynamics in both experimental EEG data and simulation. Our approach unified two distinct modes of spatiotemporal coding dynamics observed near the boundary of awareness – an early posterior dynamic code with later stable code under visibility, and earlier frontal stable code under invisibility – under a single control axis tied to effective global input.

The TG analyses revealed clear distinctions in when and how information is processed under different visibility conditions. Conscious perception was associated with a late onset of stable neural code after a strong initial dynamic neural code, engaging a distributed cortical network from sensory areas to frontal cortex. Unconscious processing, by contrast, truncated this initial feedforward cascade of dynamically changing activity pattern and showed an earlier onset of frontal stable neural code. These observations resonate with elements from a few theoretical frameworks currently under intense debate^26–28^. First, the prolonged, evolving activity across posterior cortical areas in seen trials is consistent with IIT, which emphasizes extensive local processing and integration (particularly in posterior cortex) as a basis for consciousness. The emergence of a stable code after ∼200 ms in visible conditions may reflect the formation of an integrated coalitional state (a maximally informative brain state), as IIT would predict. Second, the truncated feed-forward activity in unseen trials, coupled with early frontal activation, evokes the GNWT. GNWT proposes that conscious access requires a global broadcast ignited by frontal-parietal circuits. In our data, subjectively invisible trials lacked this ignition – the frontal cortex did activate briefly and early but did not trigger widespread persistent activity – which is in line with GNWT’s notion of a missing global workspace entry for unconscious stimuli. Instead, the top-down signals from frontal to sensory areas observed in invisible trials could correspond to attempts at ignition that remain too weak or too fleeting to produce awareness. In SI condition with trial-to-trial variability of conscious access, however, we cannot rule out the possibility that the later stable neural codes in sensory regions represent the rescued stimulus information that was weakly encoded in earlier stages. Thus, rather than contradicting each other, IIT and GNWT each capture different aspects of our findings: conscious perception near threshold requires both strong local integration (posterior cortical processing) and global broadcasting (frontoparietal feedback), and a deficit in either dimension (insufficient bottom-up drive or lack of sustained integration) results in non-conscious processing. We also note that stable neural codes observed in all ROIs regardless of whether stimulus was visible or not fits Higher-Order Theory (HOT) views in which robust first-order sensory representations can persist without awareness unless re-represented by higher-order states (often linked to prefrontal metacognitive systems)^29^.

Our connectome-based biophysical model bridged these macroscopic visibility-dependent spatiotemporal coding dynamics to mesoscopic circuit-level global input gain parameters, which is consistent with one previous computational fMRI study demonstrating that neural gain parameter could alter brain dynamics and transit the brain from a segregated to an integrated neural architecture, emphasizing the importance of information integration^30^. They also showed that the brain exhibited maximal communicability and temporal topological variability at the critical boundary between segregated and integrated states. Critically, they also showed that the effect of neural gain was most prominent in high-degree fronto-parietal network hubs. In current study, the *γ* _*PY*_ /*γ*_*VIP*_/*γ* _*PV*_ parameters effectively quantified the balance of feed-forward vs. feedback influence in each cortical area. In conscious trials, high *γ* _*PY*_ /*γ*_*VIP*_/*γ* _*PV*_ in sensory regions and low *γ* _*PY*_ /*γ*_*VIP*_/*γ* _*PV*_ in frontal cortex indicate that the network was configured to favor bottom-up ignition-like flow of information into frontal regions. In unconscious trials, the network flipped to a mode favoring top-down or local processing in frontal areas (high frontal *γ* _*PY*_ /*γ*_*VIP*_/*γ* _*PV*_, meaning little incoming drive) and greater influence of feedback in sensory areas (low *γ* _*PY*_ /*γ*_*VIP*_/*γ* _*PV*_, meaning those areas were strongly driven by inputs from elsewhere). This differential configuration aligns with the idea that in unseen trials the frontal cortex is not effectively engaged by sensory input (no ignition), and any processing in frontal areas may fail to entrain the rest of the brain, whereas in seen trials frontal areas receive strong inputs and thereby participate in a global cortical coalition. Again, as there is a trial-to-trial variability of conscious access among invisible trials, the fact that sensory regions received strong signals from other ROIs and exhibited late stable neural codes may indicate that weakly encoded stimulus information was rescued through frontal activation. A previous behavioral study reported that postcued attention could reach into the past and bring a stimulus too faint to see into consciousness by acting on the memory trace of the stimulus that disappeared before being attended^31^. Consistent with recent research suggesting that such temporally flexible conscious perception can be implemented through long-distance information sharing^15–17,19,32^. Notably, these circuit-level differences emerged even when objective performance was near chance, highlighting that subjective visibility per se correlates with a switch in neural processing regime. By linking such model parameters to theoretical concepts, we move closer to a unified account of conscious access that spans multiple scales of brain organization.

In summary, our study illustrates that near the threshold of awareness, the brain tiptoes between two modes of operation – one that amplifies a sensory event into consciousness, and one that lets it fade into unconsciousness. Using a synergistic approach of source-imaged EEG decoding and biophysical modeling, we identified a specific circuit parameter (global input gain) that differentiates these modes, offering a candidate mechanism for gating conscious access. These findings advance our understanding of the neural correlates of consciousness by providing not just correlations, but a plausible mechanistic bottom-up explanation for how subjective experience can flicker in and out without forcing the observed phenomena into existing theoretical frameworks of consciousness. Future research should build on this multi-modal framework, for instance by incorporating subcortical structures (thalamo-cortical loops) into the models, to further elucidate the pathways by which brain networks orchestrate the emergence of consciousness. Our results underscore that bridging empirical data with theoretical models is a powerful strategy to unravel the enduring mystery of conscious awareness.

## Methods

### Observers

A total of 32 human observers (18 males, 14 females; mean age = 25 ± 4.3 years) participated in this study. Data from five observers were excluded due to excessive eye movements, leaving 27 participants for analysis. EEG and behavioral data were collected from all 27 participants (21 completed two sessions, 6 completed one session; see EEG experiment). Behavior data and sensor level EEG were analyzed for 27 participants (Fig. 2 – 3).

Eye-closed structural MRI/DTI data were acquired from 21 healthy individuals (12 males; mean age, 24 years; SD, 4.2 years) using a Siemens Magnetom Prisma 3 Tesla scanner with a 64-channel head coil located at Institute for Basic Science, Sungkyunkwan University, Suwon, South Korea. Among 21 participants, 18 observers completed two sessions of EEG experiment and 3 observers completed one session of EEG. Source level EEG data were analyzed for these 21 participants (Fig. 4 – 6; Extended Data Fig. 2 – 9).

All observers had normal or corrected-to-normal visual acuity, gave informed written consent to participate as paid volunteers, and were tested individually in a dark room. The study was approved by the Institutional Review Board of the Korea National Institute for Bioethics Policy and SungKyunKwan University.

### Stimulus

Visual stimuli were generated and presented using Psychophysics Toolbox along with custom scripts written in MATLAB. A 19-inch display CRT monitor (ViewSonic PF817) was set to a refresh rate of 100 Hz and a resolution of 1024 x 768 pixels. Stimulus presentation was synchronized with the 10 ms vertical refresh cycle of the screen. The CRT monitor gamma tables were adjusted to ensure response linearity and a constant mean luminance of 59 cd/m ^2^. The target, mask, and probe stimuli had a fixed size of 16° of visual angle. Participants viewed the stimuli from a distance of 60 cm in a darkened room.

### EEG experiment

Each trial began with a fixation of randomly varied duration across trials (500 to 1000 ms). A brief target Gabor patch (20 ms) was then presented and subsequently masked by a radial sinusoid (120 ms) with no inter-stimulus interval (Fig. 1). A probe Gabor patch (50 ms) was then presented, 800 ms after the onset of the target. The spatial frequency of the target and the probe was fixed to 32.5°. Next, participants were asked to make two successive decisions (within 4 s each). First, they performed a two-alternative forced choice discrimination task, in which they indicated whether the probe was tilted clockwise (right) or counter-clockwise (left) relative to the target by rotating the Griffin PowerMate knob. The rotation of the knob by more than 12° (in one of either direction) is recorded as the participants’ response. The participants were subsequently asked to report the visibility of the target (0, no visibility; 3, maximum visibility of the target, as defined by the Perceptual Awareness Scale^15,33^. They selected one of the four boxes in the screen using the same rotating knob and pressed the space bar to complete the trial. Before performing the EEG experiments, the participants were trained for nearly 30 min to ensure that they understood the task, and sensibly used the knob for all visibility ratings.

The target orientation varied randomly among six orientations (15°, 45°, 75°, 105°, 135°, and 165°), while the probe angle was consistently tilted 30° relative to the target angle, with the tilt direction (clockwise or counterclockwise) varied randomly. The contrast of the target and the probe was fixed to 50%. The contrast of the masks differed between visibility conditions and varied between 2% and 100%. Throughout the experiment, all stimuli and masks were presented within a background gray frame (RGB value 128). In the objectively visible (OV) condition, mask contrast was set to 2% to achieve clear visibility of the Gabor stimuli. In the objectively invisible (OI) condition, mask contrast was set to 100%, such that participants’ ability to discriminate between the orientations of the target and probe was expected not to be significantly different from chance. No masking efficiency experiments were carried out. In the subjective condition, mask contrast was adjusted through an adaptive 1-up 1-down staircase procedure: on the first trial of each block, mask contrast started at 40% (based on the pilot behavior experiment); following a visible response, mask contrast on the next trial was increased by 4%; following an invisible response, mask contrast was lowered by 4% (minimum contrast of 2%, maximum contrast 100% is maintained). This adjustment was intended to yield a roughly similar number of subjectively visible (SV) and subjectively invisible (SI) trials (Extended Data Fig.1).

Each participant completed 6 blocks of trials and each block consisted of 144 trials, resulting in a total of 864 trials. Of the 27 participants, 21 completed the experiment over two sessions (i.e., two days), resulting in a total of 1,728 trials per condition for each of these participants. The remaining 6 participants completed only one session, contributing 864 trials each.

### Image acquisition

The T1-weighted structural images were acquired using 1 mm^3^ isotropic voxels; TR, 2200 ms; TE, 2.28 ms; and FOV, 256 x 256 mm. T2 weighted imaging is also acquired using 1 mm ^3^ isotropic voxels; TR, 3000 ms; TE 409 ms and FOV, 256 mm by 256 mm. Diffusion MRI data were acquired using the following parameters: 2 mm^3^ voxel size; TR, 5000 ms; TE, 106 ms; FOV of 110 mm by 110 mm, and 64 directions with b = 3000 s/mm ^2^ and seven b = 0 volumes. A single b = 0 s/mm^2^ volume was obtained with the reverse-phase encoding for use in distortion correction. All participants were recruited and data acquired in accordance with local ethics committee guidelines.

### EEG preprocessing

The EEG data were collected with 128-sensor HydroCel Sensor Nets (Electrical Geodesics) at a sampling rate of 500 Hz and were preprocessed using EEGLAB toolbox in MATLAB (Mathworks Inc.). The raw EEG data were bandpass filtered from 0.3 Hz to 200 Hz. We used Artifact Subspace Reconstruction (ASR) routine^34^ to remove noisy channels, and re-referenced all data using the average as the reference^35^. We removed line noise (60, 120, and 180 Hz) using the *cleanline* EEGLAB plugin. Then, the artifactual response components induced by eye movements and blinks were identified using independent component analysis (ICA)^36^. Finally, we determined ICs that were classified as artifacts using *ADJUST* EEGLAB plugin^37^, and removed them from the data. The EEG data were epoched from -200 ms to 800 ms relative to the onset of the target stimulus and baseline-corrected by subtracting the average signal from -200 to -10ms.

### Anatomical MRI preprocessing and source imaging

T1 data processing include brain tissue extraction, automatic skull-stripping, head surface extraction, and segmentation using *recon-all* command in Freesurfer^38^. The segmented images were then loaded into Brainstorm^39^, along with head-digitized fiducials that were recorded using the Polhemus Fastrak Digitizer (Polhemus Inc., Colchester, VT, USA). Each participant’s fiducials, co-registered with the head surface, were manually checked for misalignment. Finally, the MRI data were spatially normalized to the 1 mm Montreal Neurological Institute (MNI) template^404140^.

For obtaining forward matrix, a symmetric boundary element method (BEM) surfaces were first generated from the segmented MRI with scalp/outer-skull/inner-skull conductivities respectively taken as 0.3 S/m, 0.025 S/m, and 0.3 S/m. The default cortical surface in Brainstorm with 15,000 vertices was considered. The forward matrix (of size 128 x 15,000) was then computed using the OpenMEEG adaptive integration method^41,42^. The inverse matrix (of size 15,000 x 128) was then obtained separately for the paradigms via dynamical Statistical Parametric Mapping (dSPM) approach^43^ with default Brainstorm parameters (noise covariance regularization parameter of 0.1, signal-to-noise ratio of 3, and depth weighting order 0.5). The noise covariance matrix *C* was computed by *FF*^*T*^*/N*, where the matrix *F* is the concatenation of *N*=100*z columns of 128 sensor data obtained from 100 baseline time points of z trials (z = 1728 for participants who attended two sessions and z = 864 for those who attended one session). The source waveform for each vertex was obtained by projecting the sensor waveforms onto the inverse matrix. The vertices corresponding to each of the 68-ROI DK atlas were obtained from the Brainstorm. In each ROI, the source data are organized in a three-dimensional matrix with dimensions representing vertices, trials, and time points, respectively.

### Diffusion MRI processing and connectome construction

The diffusion data were processed using MRtrix3 version 3.0 pipeline^44^. Briefly, the diffusion images for each individual were corrected for eddy-induced current distortions, susceptibility-induced distortions, inter volume head motion, outliers in the diffusion signal^45^, within-volume motion^46^, and B1 field inhomogeneities. Tractography was conducted using the fiber orientation distributions (iFOD2) algorithm implemented in MRtrix3^44^. The algorithm used fiber orientation distributions estimated for each voxel in constrained spherical deconvolution, which can improve the reconstruction of tracts in highly curved and crossing fiber regions^47^. Streamline seeds were preferentially selected from areas where streamline density was underestimated with respect to fiber density estimates from the diffusion model. To create a SC matrix, we used DK atlas and streamlines were assigned to each region in the parcellation, yielding a directed 68 x 68 anatomical connectivity matrix for each participant. In addition, we estimated the average time-delay between ROIs by calculating the average fiber length between ROIs and divided by the axon conduction velocity of 3.9 m/s.

### Analyses

#### Behavior data

We computed the signal detection sensitivity index, *d’*, to quantify the perceptual discriminability between target and probe orientations. Specifically, responses categorized as left in trials were identified as hits when the probe was presented to the left of the target. Conversely, these responses were considered false alarms in trials where the probe appeared to the right of the target. To address extreme hit and false alarm rates (0 or 1), we applied a correction ^48^. That is, rates of 0 were converted to 1/(2*N*), and rates of 1 to 1-1/(2*N*), where *N* is the number of trials. The sensitivity index was then calculated as *d’* = Φ^-1^(hit rate) - Φ^-1^(false alarm rate), where Φ(·) is the cumulative distribution function (CDF) of the standard normal distribution.

### Multivariate pattern decoding

For decoding, the EEG data were smoothed using a 10-point moving-average filter (20 ms sliding window). The EEG data were organized in a three-dimensional matrix with dimensions representing electrodes, trials, and time points, respectively. Multivariate pattern decoding was performed participant-wise by employing a linear discriminant analysis (LDA)-based classifier with default parameters implemented in the python scikit-learn module^49^. In particular, at each time point *t*, EEG sensor/source measurements were concatenated to form an evoked pattern vector. The pattern vectors for each pairwise combination of target orientations were discriminated using LDA. The decoder performance was measured using the area under the curve (AUC) of the receiver operating characteristic curve^44^. The averaged AUC value from the decoding of fifteen pairwise combinations of target orientations was regarded as the decoder’s performance at time point *t*. The decoder was trained with OV condition and was tested with all conditions. For testing the OV condition, a 5-fold cross validation procedure was employed.

### Temporal generalization (TG)

To explore the temporal generalizability of neural representations, the multivariate pattern decoder trained at a single time point *t*_*1*_ was tested at time point *t*_*2*_ resulting in an average AUC for the pair (*t*_*1*_, *t*_*2*_). For all pairs of training and testing time points, the average AUC is computed and organized into a matrix TG(*t*_*i*_, *t*_*j*_), where *i, j* = *1,2,3*,…*499*.

### Stable/dynamic coding index

The stable and dynamic indices^5^ were used to quantify the extent of stable and dynamic coding from the TG matrix, respectively. To assess dynamic index (DI), we tested (permutation t-test) whether an off-diagonal element TG(*t*_1_,*t*_2_) was significantly smaller than both corresponding on-diagonal elements TG(*t*_1_,*t*_1_) *and* TG(*t*_2_,*t*_2_) in the TG matrix by the conjunction:

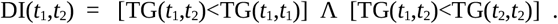

To determine stable index (SI), we tested whether the off-diagonal element TG(*t*_1_,*t*_2_) was significantly greater than 0.5 (chance level) while simultaneously not being significantly smaller than the corresponding on-diagonal elements TG(*t*_1_,*t*_1_) *and* TG(*t*_2_,*t*_2_), that is,

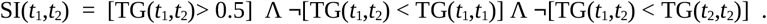

The dynamics, DI(t) and SI(t), were then obtained from the the binary matrices DI(*t*_1_,*t*_2_) and SI(*t*_1_,*t*_2_) by computing the proportion of ones (significant elements) over a 250 ms square window centered around the diagonal time point *t*. To account for the temporal smearing caused by 20 ms moving-average filter, the elements within ±10 ms of the diagonal axis were excluded while computing the proportion. A stable or dynamic index of 1 signifies a fully stable or dynamic coding, while an index of 0 indicates no dynamicity or stability in the neural representation.

### Cluster-permutation test and bootstrapping for TG matrices

The significant contours of the TG matrices were obtained by using the cluster-based permutation test^50^ that intrinsically corrects for multiple comparison issues. The standard error of the stable/dynamic index was calculated using the bootstrapping method. Specifically, a bootstrapped sample (across participants) was drawn with replacement, and its mean, referred to as the mean index, was calculated. This process was repeated 10,000 times, resulting in 10,000 mean values, from which the standard error was computed by computing the standard deviation of the 10,000 mean values.

### Cortical Microcircuitry: Fitting and Parameter Estimation

#### Single-ROI

A five-cell canonical mesoscopic model^11,12^ was implemented to simulate the source EEG waveforms for each ROI in the atlas. A cell in the mesoscopic model does not correspond to a single neuron, but rather indicates the aggregate behavior of a neural population. The model represents a local cortical microcircuitry (for a patch of cortex) consisting of two excitatory cells namely pyramidal (PY) and supplementary PY (SPY) and three inhibitory interneurons namely parvalbumin positive (PV) basket cells, somatostatin positive (SST) interneurons and vasoactive intestinal peptide (VIP) expressing interneurons. Standard local structural connections between these cells were adopted^12,23^ (Extended Data Table 1). This microcircuitry allows for a more biologically plausible representation of source local field potentials.

#### Multi-ROI and parameters

In our model, the PY cell establishes the primary long-range projections as in the real cortical networks. The PY cell in each ROI forms long-range connections with all five cell types in other ROIs, creating a total of 67 distinct long-range PY projections per ROI. The strength of these connections is determined by an anatomical connectivity matrix. The efficacy of these cortico-cortico connections to each cell is modulated by a global input gain parameter that scales the sum of amplitudes of the 67 incoming excitatory firing rates. Consequently, each ROI has five gain parameters to estimate: *γ*_PY→PY_ (pyramidal to pyramidal simply denoted as *γ*_PY_), *γ*_PY→PV_ (*γ*_PV_), *γ*_PY→VIP_ (*γ*_VIP_), *γ*_PY→SST_ (*γ*_SST_) and *γ*_PY→SPY_ (*γ*_SPY_). Additionally, the time constant of the PV cell τ_PV_ in each ROI was set as a free local parameter, allowing for region-specific temporal dynamics. This configuration enables the model to capture both the long-range interactions between ROIs and the local dynamics within each ROI, providing a comprehensive framework to evaluate the theories of consciousness. The integration of long-range cortico-cortical projections with local microcircuit interactions allows for the exploration of how global brain states emerge from distributed neural activity patterns.

#### Dynamics

The model’s dynamics were developed using a synapse-driven approach^51–54^. which is one methodological contribution of our paper. In this method, the role of a synapse— transforming postsynaptic potentials (PSP) into an average firing rate—was represented by a second-order non-linear differential equation for each cell. To facilitate solving this equation using state-space matrices, it was transformed into two simpler first-order ordinary differential equations (ODE). As a result, two state variables per synapse were created in the state-space representation. The two state variables are postsynaptic potential of a cell and its time derivative. The following first-order ODEs describe the PSP for each synapse for an ROI-*m* in the model, *m* = 1,2,3,…,68.

1. PY cell: PSP due to SPY cell:

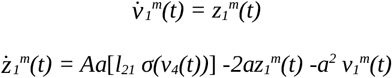 PSP due to PV:

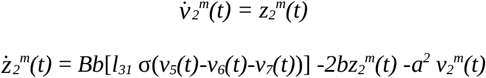 PSP due to SST:

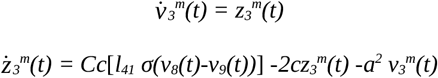
2. SPY cell: PSP due to pyramidal

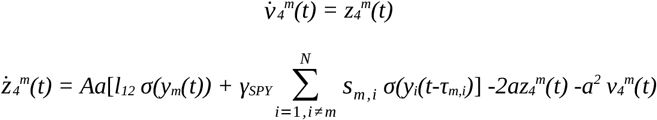
3. PV cell: PSP due to pyramidal

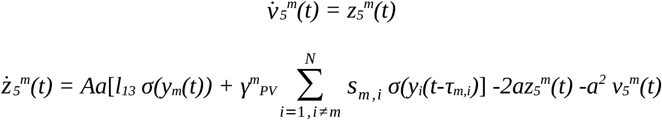 PSP due to self-inhibition in PV

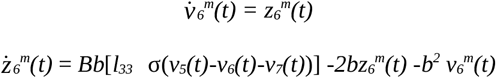 PSP due to SST

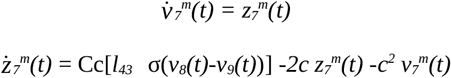
4. SST cell: PSP due to pyramidal

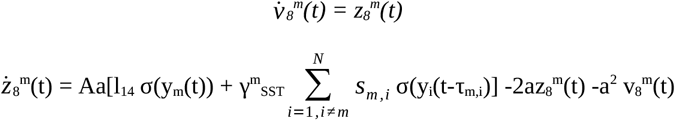 PSP due to VIP

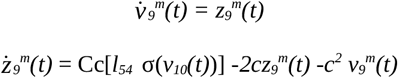
5. VIP cell PSP due to SST

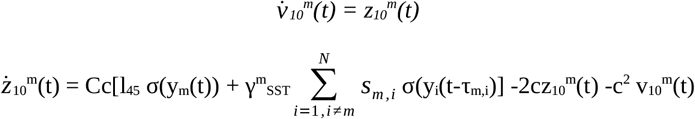 Long range pyramidal to pyramidal

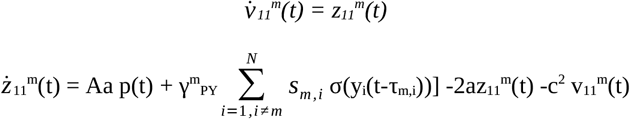

where 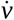 and 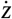 denote the first derivative of *v* and *z*, respectively. The constant A denotes the average excitatory synaptic gain and the constants B and C the average inhibitory synaptic gain, the constant *a* indicates the time-constant of excitatory post-synaptic potential, whereas the constants *b* and *c* are the time-constants of inhibitory post-synaptic potential. The cortical microcircuit model thus comprises 11 synapses (10 local and 1 global), which generates 22 state variables (shown above) represented by ODEs to capture the comprehensive dynamics of neural interactions. Six additional parameters were represented as first-order trivial dynamics. Consequently, 28 state variables in total were required to be estimated in each ROI. Except for the τ_PV_, the model’s local connectivity parameters (Extended Data Table 1), synaptic gain and time constants (Extended Data Table 2) were set to known standard values^12,23^. In the above model σ(v) = e_0_ /(1+ exp(-r (v-v_0_))) represents the sigmoidal function that characterizes the saturation and threshold effects taking place at the soma level. In that function, e _0_, r and v_0_ are the parameters that corresponds to maximum firing rate, slope of the curve and mean firing rate threshold (half-maximal response), respectively. We used the typical values of e_0_ = 2.5 Hz, r = 0.56 mV^-1^ and v_o_ = 10 mV in this paper. The s_m,i_ represents the anatomical connectivity value from ROI-*m* to ROI-*i* and τ_m,i_, estimated time-delay ROI-*m* to ROI-*i*. The aggregate postsynaptic dendritic potential to the PY cell, 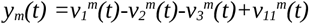 represent the simulated source waveform^49^ that forms the observation for the dynamical model.

The above ODEs may be written in the continuous-state space form as

State equation:

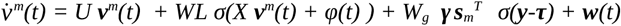

Measurement equation:

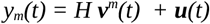

where the state vector 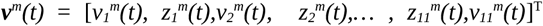, and 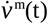 its derivative. The matrix U is a block-diagonal with entries block-diag(*M*_*1*_, *M*_*2*,_ *M*_*3*,_ *M*_*1*,_ *M*_*1*,_ *M*_*2*,_ *M*_*3*,_ *M*_*1*,_ *M*_*3*,_ *M*_*3*,_ *M*_*1*_); where 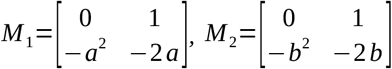, *and* 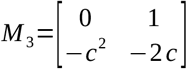. The local connectivity matrix *L=diag(0,l*_*21*_,*0,l*_*31*_,*0,l*_*41*_,*0,l*_*12*_,*0,l*_*13*_,*0,1*_*33*_,*0,l*_*43*_,*0,l*_*14*_,*0,l*_*54*_,*0,l*_*45*_,*0,0)* is diagonal and the matrix *X* is sign matrix that specifies the sign of the excitatory and inhibitory synapse. The above state and measurement equation used by the estimator (below) for determining the free parameter values.

#### Estimator

Unscented Kalman Filter (UKF) estimator^20^ was employed to calculate the state and model parameters, with one UKF utilized per ROI. The UKF in each ROI sequentially fits the source waveform in that ROI, while receiving long-range projections from the UKF in other ROIs. Unscented transform used in the estimator was set with standard parameter values^22^, namely α=1e^-3^, β=2, and κ=0. Merwe’s scaled sigma point algorithm^21^ was used to compute the sigma points of the transformation (see for the details of the unscented transformation and the UKF).

#### Fitting

With one UKF per ROI, whole brain is represented as 68 coupled UKFs. The coupled UKFs are fed with data whose dimension is 68 x time-points. After sequential fitting, the fitted data of the same dimension and six estimated parameters were obtained. To extract the parameters vertex-wise (for each trial), we carried out a vertex-wise fitting procedure as follows: vertex data in one target ROI (vertices x time-points) is kept intact and waveforms in other ROI are averaged across vertex resulting in 67 x time-points data. Now, by taking one vertex waveform at a time in the target ROI and the averaged waveforms from other ROI, we fit the coupled UKF. This procedure is repeated for all the vertices in the target ROI. The entire procedure is then repeated for each of the 68 ROIs participant-wise. Through this vertex-wise fitting procedure, simulated source data are organized into a three-dimensional matrix with dimensions representing vertices, trials, and time points. This simulated source data is subsequently used for decoding analyses (Fig. 6b). Each extracted parameter waveforms were fitted with exponential waveforms of the form *C+A*(1-exp(-Bt))* to extract the exact parameter value and the convergence time (Extended Data Fig. 8).

#### FDR correction

To assess the statistical significance of whole-brain vertex-wise analyses, we employed a cluster-based permutation approach with False Discovery Rate (FDR) correction. For each of the 15,000 vertices across 21 participants, we computed the observed test *t*-statistic (*t*-statistic). Clusters were formed by grouping adjacent vertices that exceeded a predefined threshold (p < 0.05, uncorrected). We then calculated cluster-level statistics (sum of t-values within each cluster). A permutation procedure was implemented, randomly shuffling participant labels 10,000 times, and repeating the clustering process for each permutation. The observed cluster-level statistics were compared against the distribution of maximum cluster-level statistics from permutations to derive p-values. Finally, we applied FDR correction to the cluster-level p-values, sorting them in ascending order and calculating the FDR threshold with a desired FDR level (q=0.05). Clusters with FDR-corrected p-values below the significance threshold were considered statistically significant, allowing us to identify brain surface clusters while controlling for multiple comparisons.

## Data and Code availability

The data and code supporting the findings of this study are available at https://osf.io/g8s2m. Access is currently restricted, but will be openly released upon publication. The archived repository contains the raw EEG and MRI/DTI data, forward and inverse matrices for source reconstruction, as well as the Python scripts used for decoding analyses and all results presented in this study.

## Extended Data

**Extended Data Fig 1.**
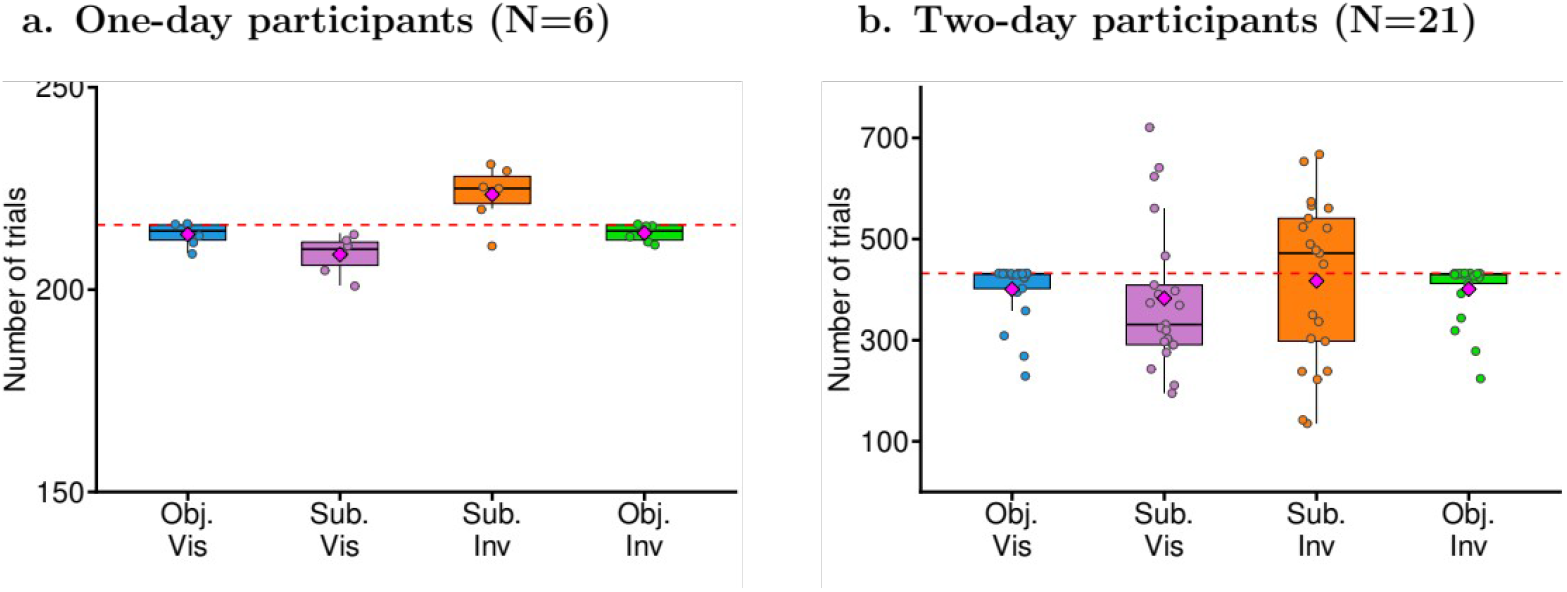
Trial distribution across conditions after post-artifact rejection. **a**. Trial counts for one-day participants (*N*=6) across visibility conditions. In objective visible/invisible conditions, 216 trials were presented (red dashed lines). Subjective visible trials were decreased while subjective invisible trials increased, reflecting individual differences. (OV: 213 ± 3; SV: 208 ± 5; SI: 224 ± 7; OI: 214 ± 2) **b**. Trial counts for two-day participants (*N*=21), with 432 trials presented in objective visible/invisible conditions (red dashed lines). Overall, the balanced distribution of trials across conditions/paradigms (OV: 401 ± 59; SV: 383 ± 144; SI: 417 ± 163; OI: 401 ± 59) mitigates classification biases due to imbalanced trials across conditions. This ensured us robust statistical comparisons of neural representations across varying levels of conscious access.

**Extended Data Fig 2.**
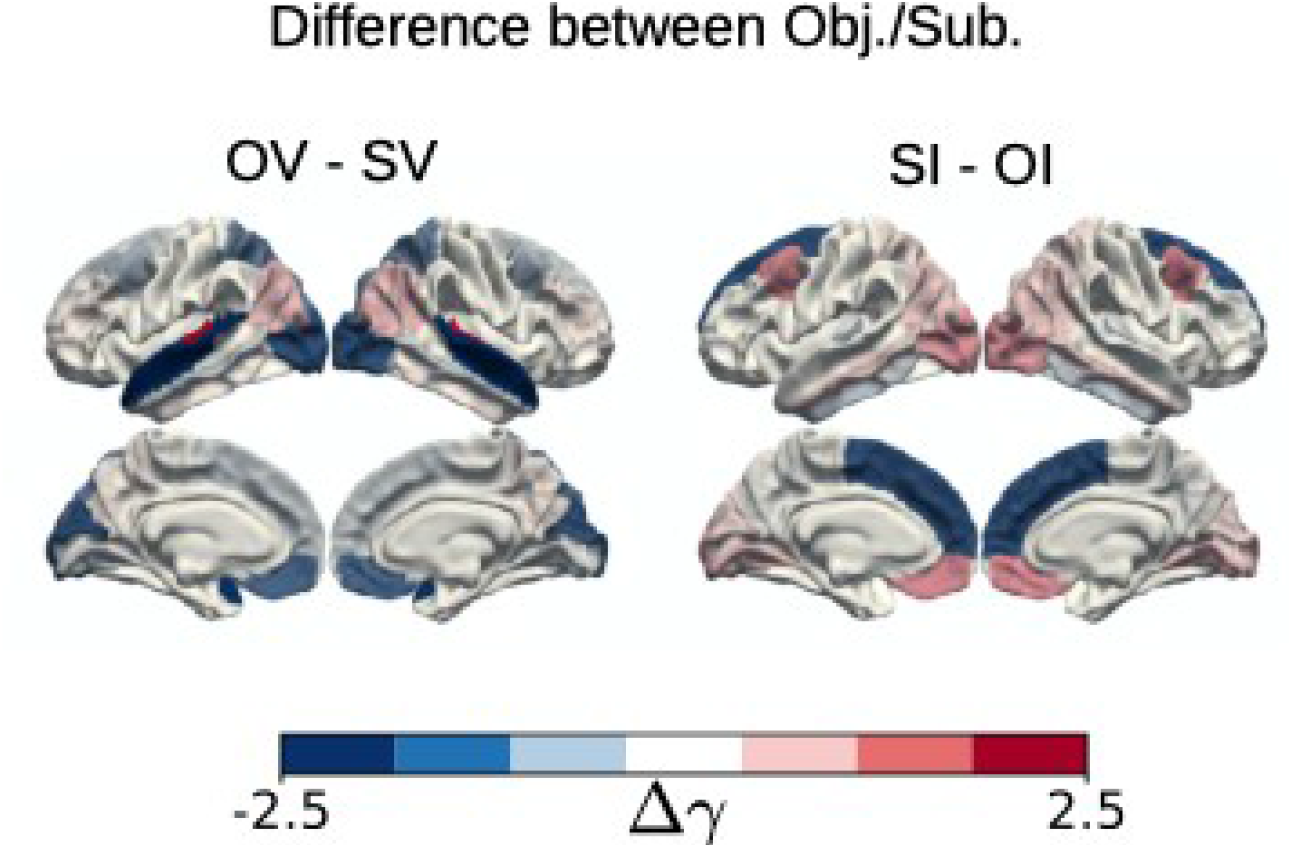
*γ* _*PY*_ differences in OV – SV and SI – OI maps (from Fig. 5a). OV condition shows higher *γ* _*PY*_ values in parietal and temporal regions, indicating enhanced local processing in those areas. SV condition displays elevated values in visual, parietal, temporal, and select frontal areas, suggesting a broader network engagement during conscious perception. SI condition exhibits positive *γ* _*PY*_ values across visual, parietal, temporal, and frontal cortices, potentially suggesting altered information flow for the subliminal stimuli. OI condition shows negative values primarily in frontal cortices, which may indicate reduced local processing and increased susceptibility to top-down modulation during unconscious stimulus processing.

**Extended Data Fig 3.**
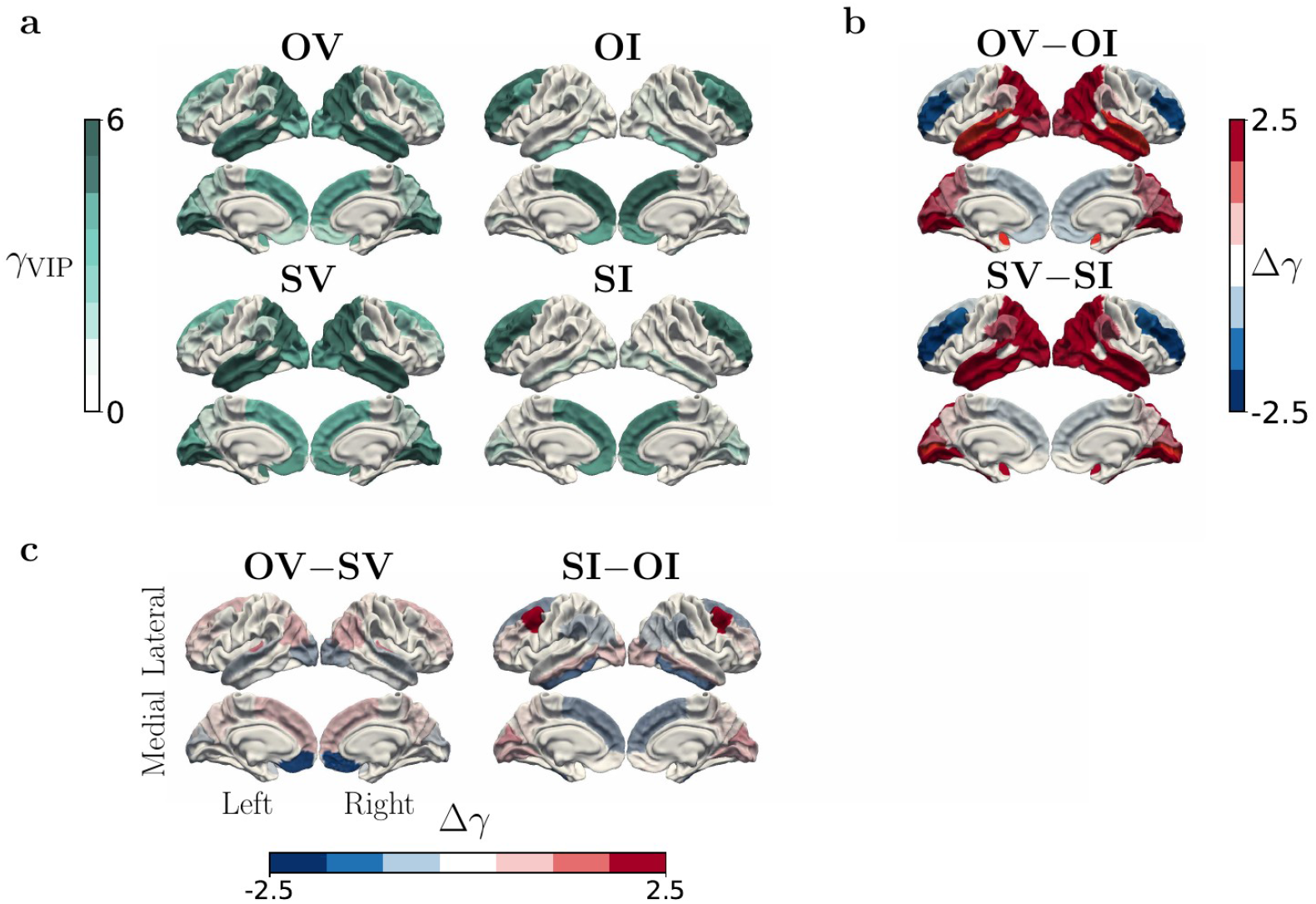
Surface maps of VIP global input gain (γ_VIP_) distribution reveals differential engagement during varying levels of conscious access.

**Extended Data Fig 4.**
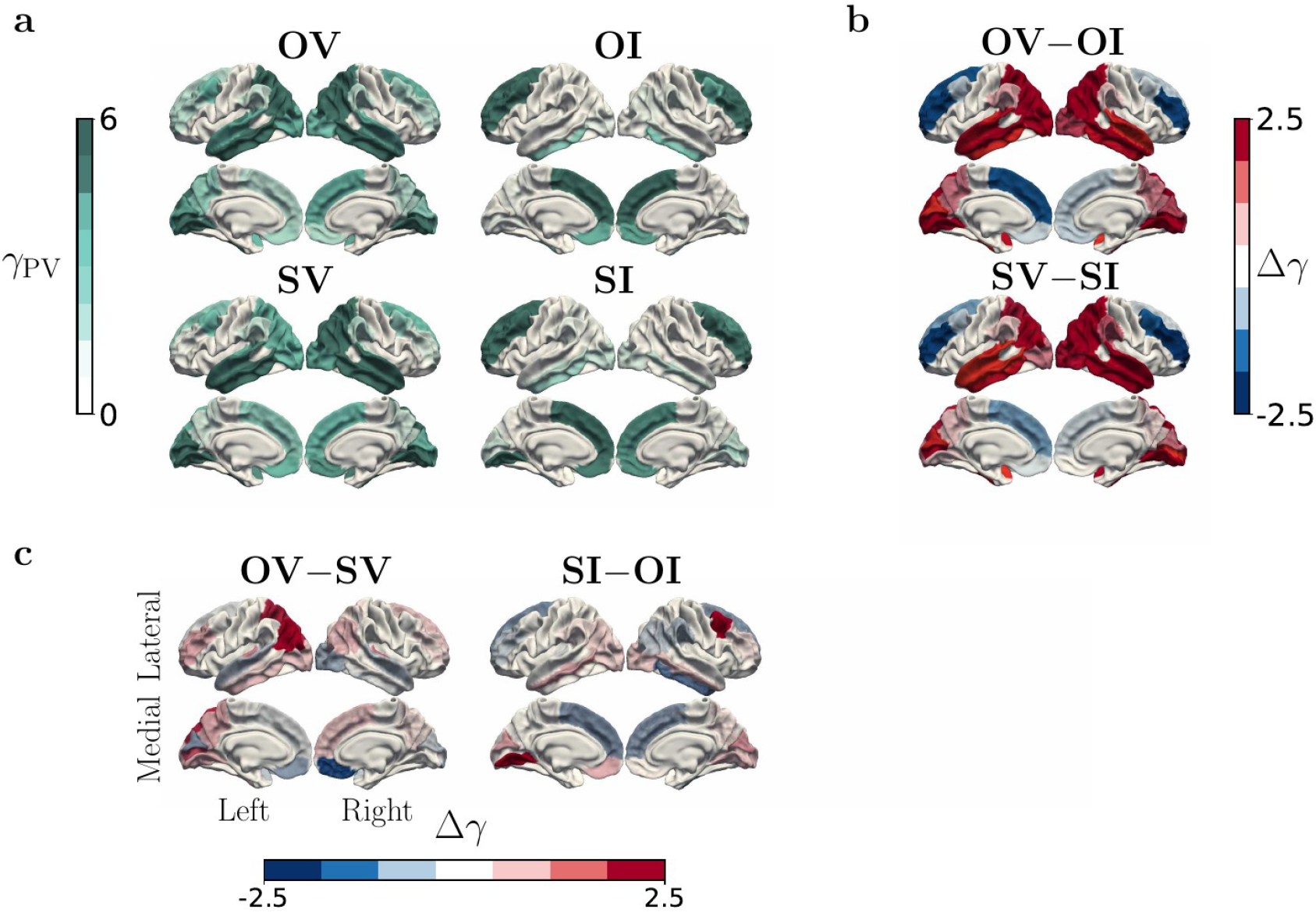
Surface maps of PV global input gain (γ_PV_) distribution reveals differential engagement during varying levels of conscious access.

**Extended Data Fig 5.**
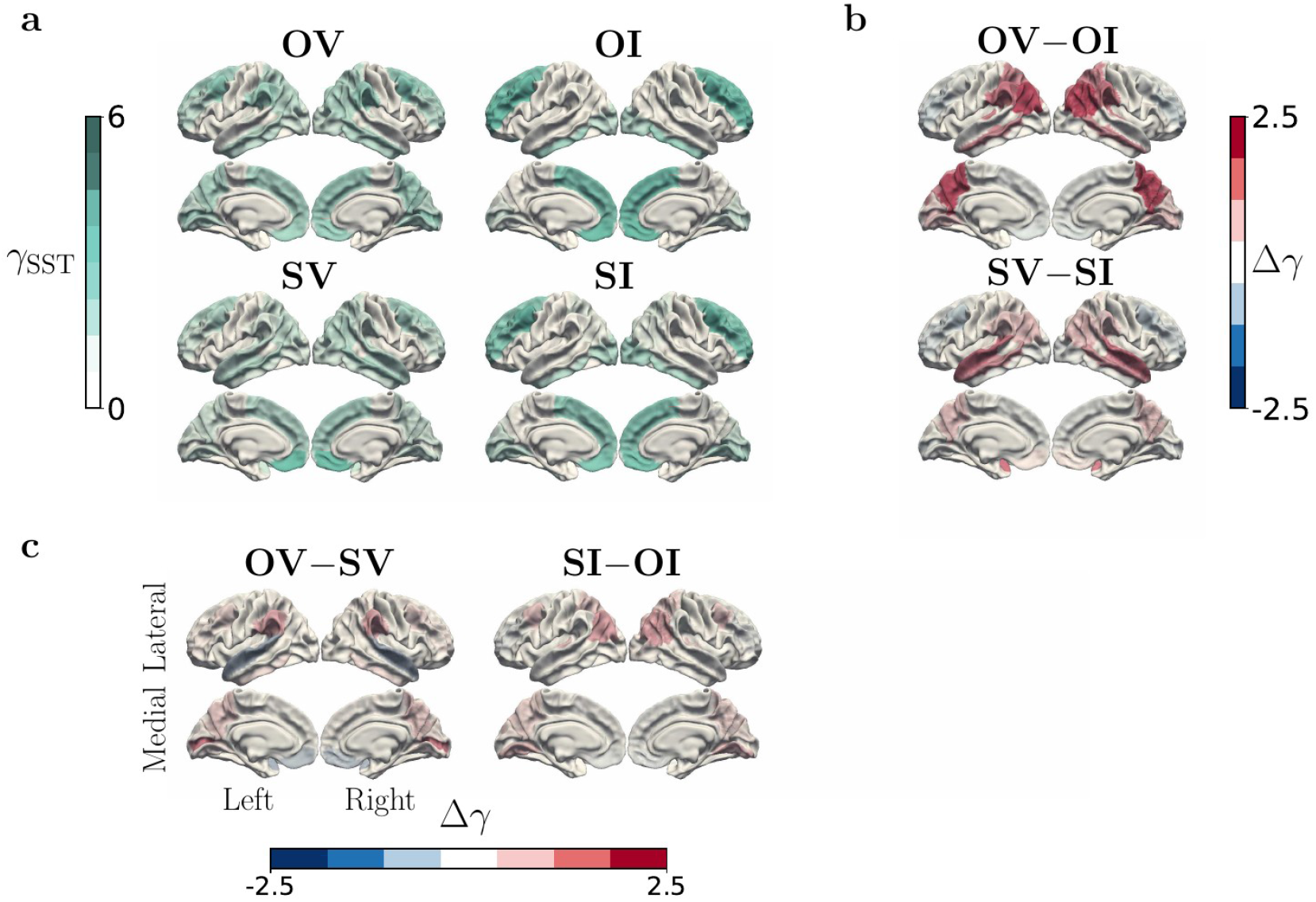
Surface maps of SST cell global input gain (γ_SST_) distribution reveals differential engagement during varying levels of conscious access.

**Extended Data Fig 6.**
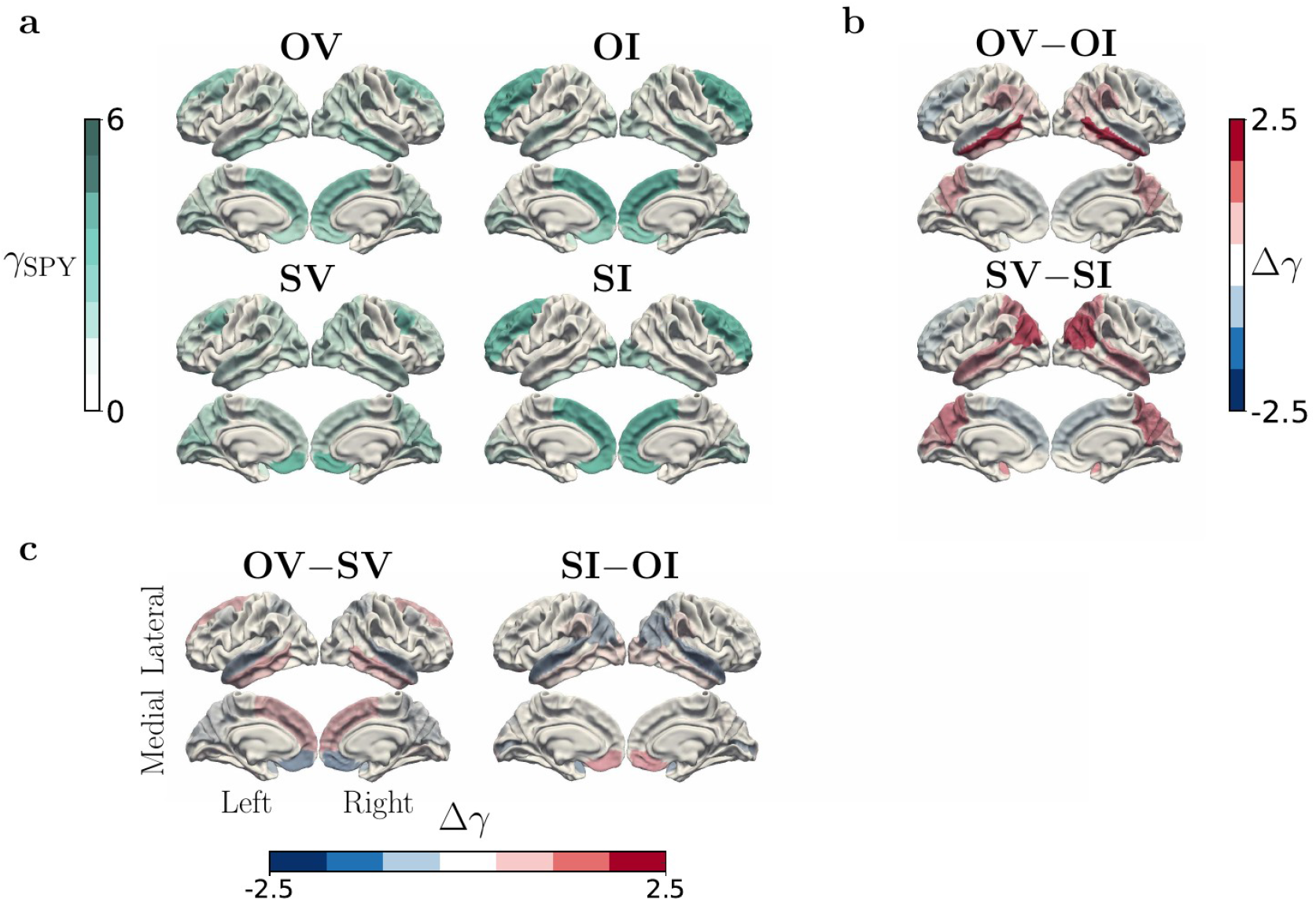
Surface maps of supplementary pyramidal cell global input gain (γ_SPY_) distribution reveals differential engagement during varying levels of conscious access.

**Extended Data Fig 7.**
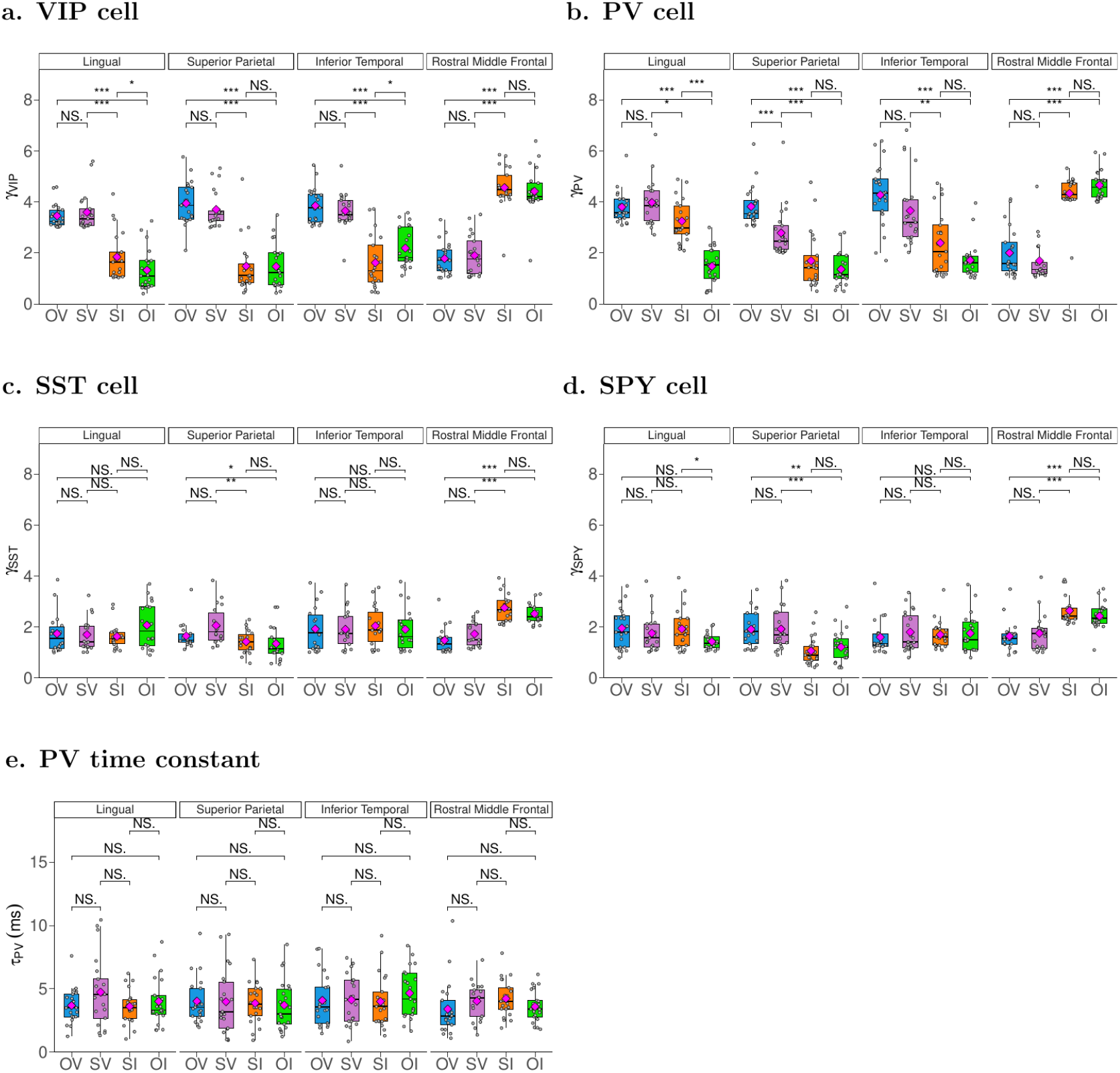
**Global input gain** of **a**. VIP **b**. PV **c**. SST **d**. SPY neuronal populations for four representative region of interest (ROI). The VIP and PV populations exhibited input gain patterns similar to those observed in the pyramidal population (Fig. 5c) across the ROIs. In contrast, the SST and SPY populations did not display consistent patterns of change in input gain. The similar input gain patterns of VIP and PV populations compared to pyramidal neurons may contribute to the establishment of distinct cortical regimes, such as the high-gain state at low mask contrast (visible) and the low-gain state at high contrast (invisible) **e**. The local PV time constant showed no significant differences among the ROIs. This suggests that PV population provided consistent inhibition (no regional variability) throughout the cortex and thus, consistent sensory integration windows. However, the input gain of the PV significantly varies across the ROIs/conditions (Extended Data Fig. 4), suggesting flexible gain control of sensory representations while maintaining a consistent temporal integration.

**Extended Data Fig 8.**
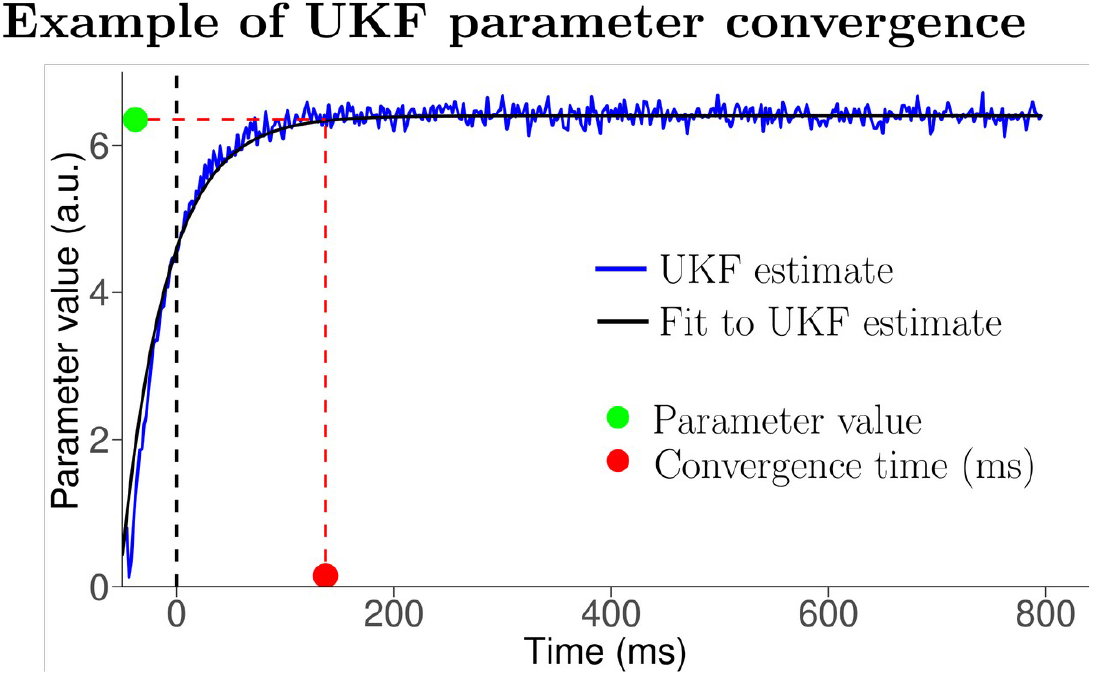
Example of parameter convergence time and parameter value determination for one vertex source data. An exponential curve is fitted to the UKF estimates to determine both the converged parameter value and convergence time.

**Extended Data Fig 9.**
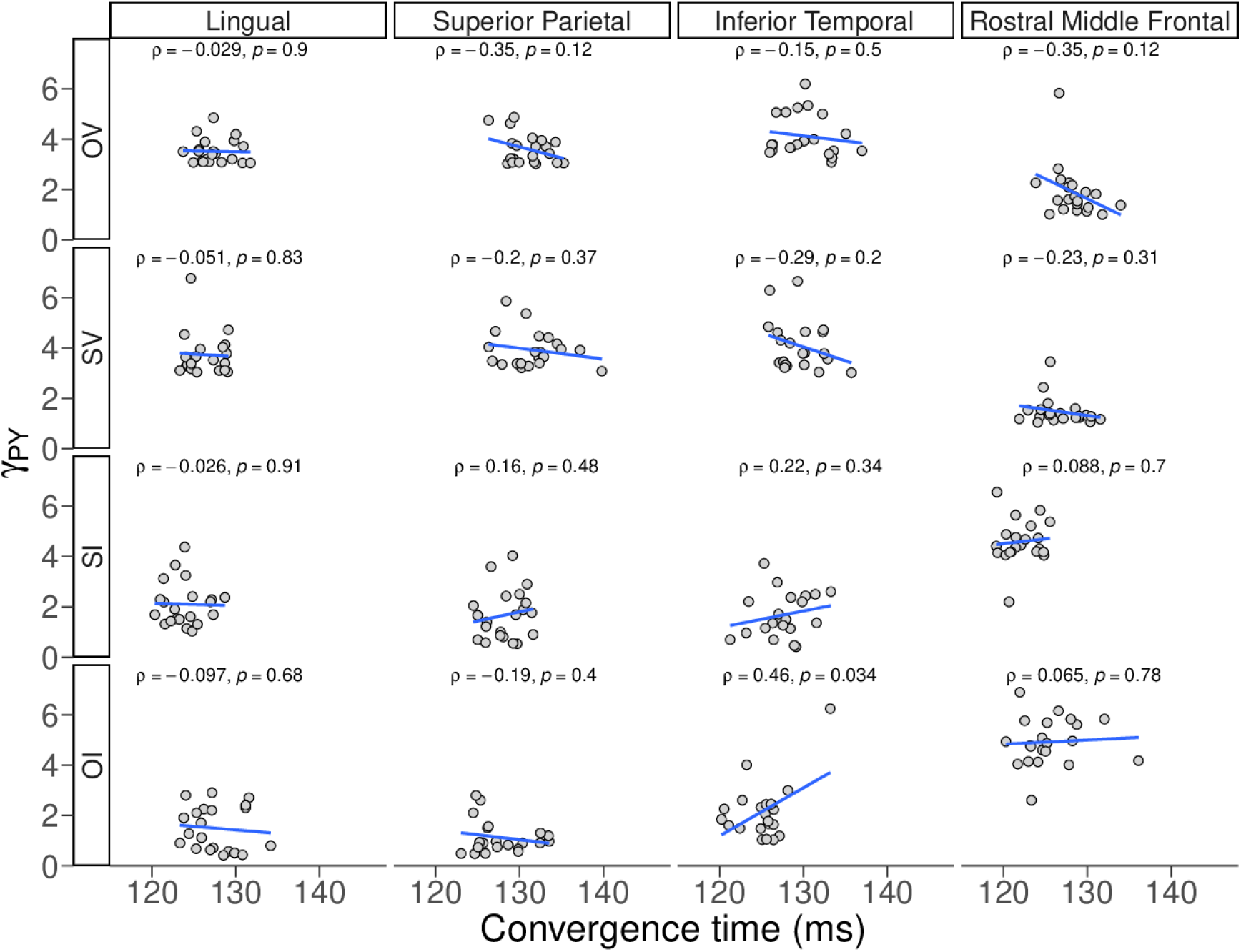
Correlation patterns (blue lines) between the magnitude of the γ_PY_ (Fig. 5c) its convergence time (Fig. 5d) reveal distinct patterns for visible and invisible conditions. This figure illustrates the correlation pattern (blue line) between pyramidal cell global input gain (γ_PY_) and the convergence time across four key regions of interest (ROIs): Lingual (L), Superior parietal (SP), Inferior temporal (IT), and Rostral middle frontal (RMF). The patterns were shown for the four levels of conscious access: objective visible (OV), subjective visible (SV), subjective invisible (SI), and objective invisible (OI). In the visible conditions (OV/SV), γ_PY_ gradually decreased along the processing hierarchy from L to SP to IT to RMF. Convergence times remained moderate (120 to 140 ms) across these regions. In the invisible conditions (SI/OI), γ_PY_ values showed a gradually increasing pattern from L to SP to IT to RMF. Convergence times were the least in the SI condition. This pattern was a reversal compared to the visibility conditions. While the convergence time of parameters in the unscented Kalman filter estimator may not directly indicate the earlier onset of stable coding in prefrontal areas in the SI condition in particular. However, its correlation with the magnitude of γ_PY_ changes gradually from posterior to prefrontal areas, hinting at a potential link to the temporal dynamics of information processing.

**Extended Data Table 1.** Local circuit connectivity constants (*l*_*ij*_s in the model equations)

| From/To | PY | SPY | PV | SST | VIP |
| --- | --- | --- | --- | --- | --- |
| PY | 0 | 135 | 30 | 33.75 | 0 |
| SPY | 108 | 0 | 0 | 0 | 0 |
| PV | 550 | 0 | 100 | 0 | 0 |
| SST | 33.7 | 0 | 13.5 | 0 | 20 |
| VIP | 0 | 0 | 0 | 200 | 0 |

**Extended Data Table 2.** Biophysical parameters of the model.

| Cell type | Parameter | Value | Description |
| --- | --- | --- | --- |
| **PY & SPY** | A | 4 mv | Average excitatory synaptic gain |
|  | a | 100 s<sup>-1</sup> | Time-constant of excitatory post-synaptic potential |
| **PV**<br>**(fast GABA)** | B | -20 mv | Average inhibitory synaptic gain |
| | b | 1/$\tau_{PV}$ s<sup>-1</sup><br>(free local parameter) | Time-constant of inhibitory post-synaptic potential |
| **SST & VIP**<br>**(slow GABA)** | C | -30 mv | Average inhibitory synaptic gain |
| | c | $20\text{ s}^{-1}$ | Time-constant of<br>inhibitory post-synaptic potential |

## Notes

### Competing Interest Statement

The authors have declared no competing interest.

